# Generation and Validation of a Lower Limb Muscle Single-Cell RNA Sequencing Data Set Identifies Pathogenic Endothelial Metabolism in Peripheral Arterial Disease

**DOI:** 10.1101/2025.09.23.678171

**Authors:** Mohamed Nofal, Suhib Alhusban, Qimei Han, Adeleigh Pressley, Samantha O’Keefe, Silvia Leanhart, Kevin Southerland, Joseph M McClung, Brian H Annex

**Affiliations:** Vascular Biology Center and Department of Medicine, Medical College of Georgia at Augusta University, Augusta, GA, USA; Department of Surgery, Duke University School of Medicine, Durham, NC; Molecular Medicine, Internal Medicine, and Vascular and Endovascular Surgery, Wake Forest University School of Medicine, Winston-Salem, NC

**Keywords:** Peripheral Arterial Disease (PAD), Chronic Limb-Threatening Ischemia (CLTI), Endothelial Cells (EC), Glycolysis, Hexosamine Biosynthetic Pathway (HBP), Single-cell RNA sequencing (scRNA-seq), supervised exercise

## Abstract

**Background:** Peripheral arterial disease (PAD) results from atherosclerotic occlusion(s) in leg arteries. Chronic limb-threatening ischemia (CLTI) is the most severe form of PAD. Patients with CLTI suffer from rest pain, ulcers, or gangrene. Clinical outcomes remain poor in patients with CLTI and many investigational approaches, such as promoting angiogenesis, have failed. Understanding cell-specific vs. bulk-RNA changes within muscle offers an opportunity to better understand this disease.

**Objective:** To assess cell-specific alterations in gene and metabolism pathways in endothelial and muscle cells from leg muscle between patients with PAD vs. non-PAD controls.

**Methods:** We analyzed publicly available bulk RNA-seq leg muscle data from non-PAD controls, patients with intermittent claudication (IC) and CLTI. An emphasis was on glycolysis and the hexosamine biosynthetic pathway (HBP). Two single-cell RNA-seq datasets were integrated, cell types identified, and metabolism pathway scores calculated across identified cell types. Validations included quantitative mRNA in flow-sorted ECs, and immunofluorescence from clinically-phenotyped human samples.

**Results:** Bulk RNA-seq revealed downregulation of both glycolysis and HBP in CLTI vs. IC or control tissues. Single-cell analysis uncovered cell-type specific changes. In patients with CLTI, gene expression in ECs showed increased glycolysis and decreased HBP. In myonuclei gene expression of both pathways were reduced. Flow-sorted ECs confirmed higher glycolysis and reduced HBP in CLTI vs. controls. Immunofluorescence analysis of muscle revealed significantly lower HBP⁺ECs (9.5% vs. 31.3%, n=5/group, p=0.004) and increased glycolytic ECs (42.0% vs. 14.4%, n=7/group, p<0.001) in CLTI vs. control. The findings were independent of capillary density. In IC patients, supervised exercise vs. optimal medical care reduced ischemic muscle PFKFB3^+^ ECs (2.67% vs. 14.61%, n=11 vs. 9, p<0.001).

**Conclusions:** We generated and validated a single-cell RNA-seq data set that confirms maladaptive metabolic reprogramming in CLTI ECs. Strategies designed to alter metabolism need to account for differences between bulk vs. single-cell analyses.

## Introduction

Peripheral arterial disease (PAD) arises from atherosclerotic occlusions in the large arteries of the lower extremities that compromise tissue perfusion, and ultimately lead to skeletal muscle ischemia and functional impairment.^1,2^ The prevalence of PAD is estimated to exceed 14 million in the United States and over 200 million globally.^2^ The two primary symptomatic presentations of PAD are intermittent claudication (IC) and chronic limb-threatening ischemia (CLTI). IC is characterized by muscle discomfort in the calf or buttock during physical exertion, which subsides with rest.^2^ CLTI represents the most severe form of PAD, characterized by the presence of ischemic rest pain, non-healing ischemic ulcers, or gangrene.^2^ In the US, patients with CLTI are frequently treated with surgical and/or endovascular procedures to improve blood flow to ischemic limbs. However, many patients are not candidates for surgical or catheter-based revascularization because of excessive comorbid conditions or unfavorable vascular anatomy. In addition, often successful large vessel revascularization may be inadequate for symptom relief because of residual microvascular disease.^2,3^ Therefore, despite advancements in medical therapies across patients with cardiovascular disease in general, patients with PAD and CLTI continue to be in need for more effective therapeutic approaches.

When an arterial occlusion occurs proximal to a muscle, both the number and functional capacity of local blood vessels in the down-stream ischemic tissue are critical determinants of oxygen delivery and the clearance of metabolic waste products.^2^ Over the past two decades, numerous human studies have attempted to stimulate new blood vessel formation through therapeutic angiogenesis, yet these efforts have largely failed to translate into clinical success.^2,4–6^ Tumor blood vessels by design are structurally and functionally abnormal, resulting in leakiness and inefficient/poor oxygen delivery causing regions of hypoxia within the tumor mass.^7–9^ Tumor progression is further supported by pathological angiogenesis mediated by vascular endothelial growth factor (VEGF), which promotes glycolytic metabolism in tumor endothelial cells (ECs).^10,11^ Studies specifically investigating EC metabolism in PAD remain limited.

Cell metabolism is a complex network of interconnected pathways involving multiple metabolites whose fate is determined by the availability and activity of rate-controlling enzymes. In ECs, glycolysis is the predominant pathway for ATP production and is primarily regulated by levels of 6-phosphofructo-2-kinase/fructose-2,6-biphosphatase isoform 3 (PFKFB3), which catalyzes the production of fructose-2,6-bisphosphate, a potent allosteric activator of phosphofructokinase-1.^12,13^ In preclinical models of PAD, EC specific genetic deletion of PFKFB3 led to impaired neovascularization and compromised perfusion recovery following hindlimb ischemia (HLI).^14,15^ In contrast, in cell and murine models of PAD, EC PFKFB3 expression and glycolysis are increased.^16^ Altering glucose availability to ECs has neither beneficial nor detrimental effects.^17^ Studies that modulated, not deleted glycolysis have shown that elevated glycolytic capacity appears to have poor prognostic value, while limiting increases in glycolytic activity yields beneficial effects.^16–18^

The cell metabolism pathways of glycolysis, pentose phosphate pathway (PPP), and the hexosamine biosynthesis pathway (HBP) intersect (Figure S1).^16,18^ Glucosamine supplementation activates the HBP, leading to enhanced endothelial O-GlcNAcylation and activation of ATF4, a transcription factor known to mediate exercise-induced angiogenesis.^18,19^ The ability of HBP to function independent of changes in VEGF and glycolysis was shown by the fact that in eNOS⁻/⁻ mice, glucosamine treatment reduced tissue necrosis following HLI. Moreover, in BALB/c mice, glucosamine increased HBP activity, improved perfusion recovery, and increased capillary density.^18^ In vitro under PAD relevant conditions, glucosamine enhanced EC survival, barrier function, and tube formation without stimulating glycolysis or PPP, but rather by increasing amino acid metabolism thereby enhancing oxidative phosphorylation to generate ATP and serine biosynthesis for NADPH production.^18^ These metabolic changes were associated with reduced ROS accumulation and reduced permeability when compared to VEGF₁₆₅a-treated cells.^18^ These studies highlight that coordinated regulation of metabolic pathways is critical for effective functional vascular regeneration.

Cell type–specific metabolic changes in human skeletal muscle in the context of PAD, particularly in advanced CLTI, remain poorly defined. We sought to create an integrated single-cell RNA sequencing data set from leg muscles from humans without PAD, and patients with IC or CLTI. We sought to determine whether predictions from this data set were both present in prior published data and could be validated in human samples by performing flow cytometry–based endothelial cell isolation and immunofluorescence of predicted differential protein expression based on RNA values. Collectively, this work demonstrates that cell-specific transcriptomic data provides insights beyond those observed in bulk tissue analysis, and how maladaptive metabolic alterations arise across distinct cell types in PAD, thereby identifying new therapeutic targets to improve outcomes.

## Methods

### Input of BULK RNA Seq Data and Selective Pathway Analysis

We used publicly available bulk RNA-sequencing of the whole transcriptome shotgun sequencing (WTSS) data set derived from gastrocnemius muscle biopsies collected from three groups: non-PAD controls, patients with IC and CLTI.^20,21^

For each group, a composite pathway expression score was calculated as the average of the log-normalized expression values of genes within the selected specified pathway. These scores are reported as mean ± standard error of the mean (SEM) for each group. The specific genes used for each pathway are listed in Supplementary tables.

To maintain analytical consistency, all composite scores were computed using the same averaging method applied to log-normalized expression counts.

Visualization of individual gene expression patterns using dot plots generated in R (v4.4.2) with the *ggplot2* package. In these visualizations, each panel represents a single gene, with each dot corresponding to the expression level in one individual sample.

### Generation of a Single-cell RNA Sequencing (scRNA-seq) Dataset from Lower Limb Muscle from Control, IC, and CLTI

Single-cell RNA sequencing (scRNA-seq) data from two publicly available datasets were analyzed using Seurat (v5) in R (v4.4.2). The first dataset (GSE235143) included gastrocnemius muscle samples from non-diseased control individuals and patients with peripheral arterial disease (PAD); PAD patients who had undergone limb amputation were excluded from the study.^22^ The second dataset (GSE227077) included samples from patients with CLTI undergoing limb amputation. In this second dataset, samples were taken from both proximal and distal regions relative to the amputation site. The proximal region lacked overt tissue necrosis and was used in this analysis.^23^

From both datasets, cell barcodes were standardized to match patient metadata. Cells with low gene counts (<200 genes) or >20% mitochondrial gene content were excluded. The remaining data were log-normalized using the “LogNormalize” method (scale factor = 10,000). Highly variable genes (n = 2000) were identified using the “vst” (variance-stabilizing transformation) method, followed by scaling, principal component analysis (PCA) using the top 30 principal components, and dimensionality reduction using Uniform Manifold Approximation and Projection (UMAP). Initial integration of control and PAD samples from (GSE235143) was performed using canonical correlation analysis (CCA), and CLTI samples from (GSE227077) were subsequently integrated using Harmony for batch correction. The entire preprocessing pipeline was repeated post-integration to ensure uniformity.

Cells were clustered using Seurat’s FindNeighbors() and FindClusters() (resolution = 0.5), and annotated manually based on canonical marker gene expression visualized with FeaturePlot() and DotPlot(). Marker genes used for cell type classification included in supplementary tables.

Differential gene expression analysis was conducted using Seurat’s FindMarkers() function. While multiple pairwise comparisons were explored, the final analysis focused on CLTI vs. control.

### Validation of the sc-RNA Seq Dataset by Flow Assisted Cell Sorting from Human Muscle

Healthy adult/PAD-free control volunteers (n=5) and CLTI (n=4) patients were recruited through print advertising under approved IRB protocols. Percutaneous muscle biopsy samples were taken from the lateral gastrocnemius muscle of age-matched volunteer HAs. Muscle specimens from CLTI patients were collected form the same anatomical location, as previously described.^20^ Candidate gene expression in flow sorted cells was assessed to measure predictions from the sc-RNA data. Skeletal muscle samples were homogenized, washed, digested, and filtered to isolated cell stock for flow assisted cell sorting (FACS). Briefly, tissue was minced and centrifuged in DMEM supplemented with antibiotics before a 30-minute digestion at 37° C in 10mL Collagenase B/Dispase solution. Digested slurry is filtered through a 100µm strainer and centrifuged to clear before debris removal and resuspension in PBS supplemented with FBS. Pelleted/resuspended cellular material is then transferred to separate FACS tubes for preparation of labelled and unlabeled samples. The unstained samples were prepared by adding 100µL of the cell stock into a FACS tube with 200µL of PBS+0.2% FBS. The full stain samples were prepared by centrifugation of the remaining stock for 400xg for 7 minutes. The supernatant was then aspirated leaving approximately 90µL in the tube. After the addition of 5µL of Fc Block, the tube was incubated on ice for 15 minutes. The respective antibodies were then added and incubated in the dark for 20 minutes. Then, 3mL of 1x PBS+0.2% FBS was added, and the sample was centrifuged at 500xg for 7 minutes. The tube was decanted and brought to a final volume of 300µL with 1X PBS+0.2% FBS. Antibody capture beads were used as compensation controls. Two drops of each antibody capture bead (positive and negative) were added to a FACS tube. These beads were then equally distributed amongst three tubes to match the three fluorochrome types that were being used (APC, FITC, PE). Next, each antibody that was used with their corresponding amount was added to its respective FACS tube. They were incubated at room temperature in the dark for 15 minutes. 3mL of 1X PBS+2% FBS was added and then the sample was centrifuged at 500xg for 7 minutes. The sample was then aspirated manually, leaving 300µL in the tube. The labeled samples were incubated with CD31 (eBiosciences, #14-0319-82), CD56 (eBiosciences, 555518). Additionally, 4′,6-diamidino-2-phenylindole (DAPI; D1306, Thermo Fisher) was added to the sample to discern a live cell population. Fluorochrome controls using antibody capture beads are used as compensation controls. Endothelial cells were identified in the heterogenous cell mixture using a set around live cells positive for CD34, CD31, and SCA1. A BD FACSAria^TM^Fusion cell sorter was used for all FACS experiments. Initial gating was done to remove debris from the total cell population. This was followed by gating for single cell entities to generate a singlet population and gating a Live/Dead check, using DAPI. Lastly, a gate was set for viable cells that are CD31^+^ and CD56^-^. Studies were conducted by Dr. McClung at Wake Forest University.

### Validation of sc-RNA Seq Data by Immunofluorescence Staining of Human Muscle Sections

We used the same immunofluorescence staining protocol on histologic samples using methods recently described.^18^ For this study, new antibody combinations were used to assess endothelial PFKFB3 expression: (1) mouse anti-CD31 [JC/70A] (Abcam, Cat# ab9498) with rabbit anti-PFKFB3 [EPR12594] (Abcam, Cat# ab181861), and for endothelial O-GlcNAc expression (2) rabbit anti-CD31 [EPR3094] (Abcam, Cat# ab76533) with mouse anti-O-GlcNAc (Thermo Fisher Scientific, Cat# MA1-076). For immune cell identification, additional primary antibodies included rabbit anti-CD45 [EP322Y] (Abcam, Cat# ab40763) and rabbit anti-CD68 [EPR20545] (Abcam, Cat# ab213363). Quantification was performed as the percentage of CD31⁺ endothelial cells co-expressing PFKFB3 or O-GlcNAc across ≥5 random high-power fields per patient. Image analysis was conducted using CellProfiler, blinded to group assignment.

## Results

### Bulk vs. Single Cell RNA-Seq Measures of HBP, ATF4, and Glycolysis Across Control, IC, and CLTI Human Skeletal Muscle

We first analyzed bulk-RNA sequencing from human skeletal muscle samples to assess potential alterations in HBP and glycolytic gene expression, comparing non-PAD controls, patients with IC, and patients with CLTI.^20,21^ The HBP score was significantly reduced in CLTI (40.31 ± 1.65, mean ± SEM, *n* = 16) compared with both IC (51.13 ± 1.51, mean ± SEM, *n* = 20; *P* <0.001) and control (49.21 ± 1.85, mean ± SEM, *n* = 15; *P* = 0.007; Figure 1A). This reduction was driven by the consistent downregulation of the genes within HBP pathway, including (GFPT1, GFPT2, GNPNAT1, and OGT) used to calculate the score. Dot plots showing expression patterns of the individual genes within the HBP pathway are shown in Supplementary Figure S2.

**Figure 1.**
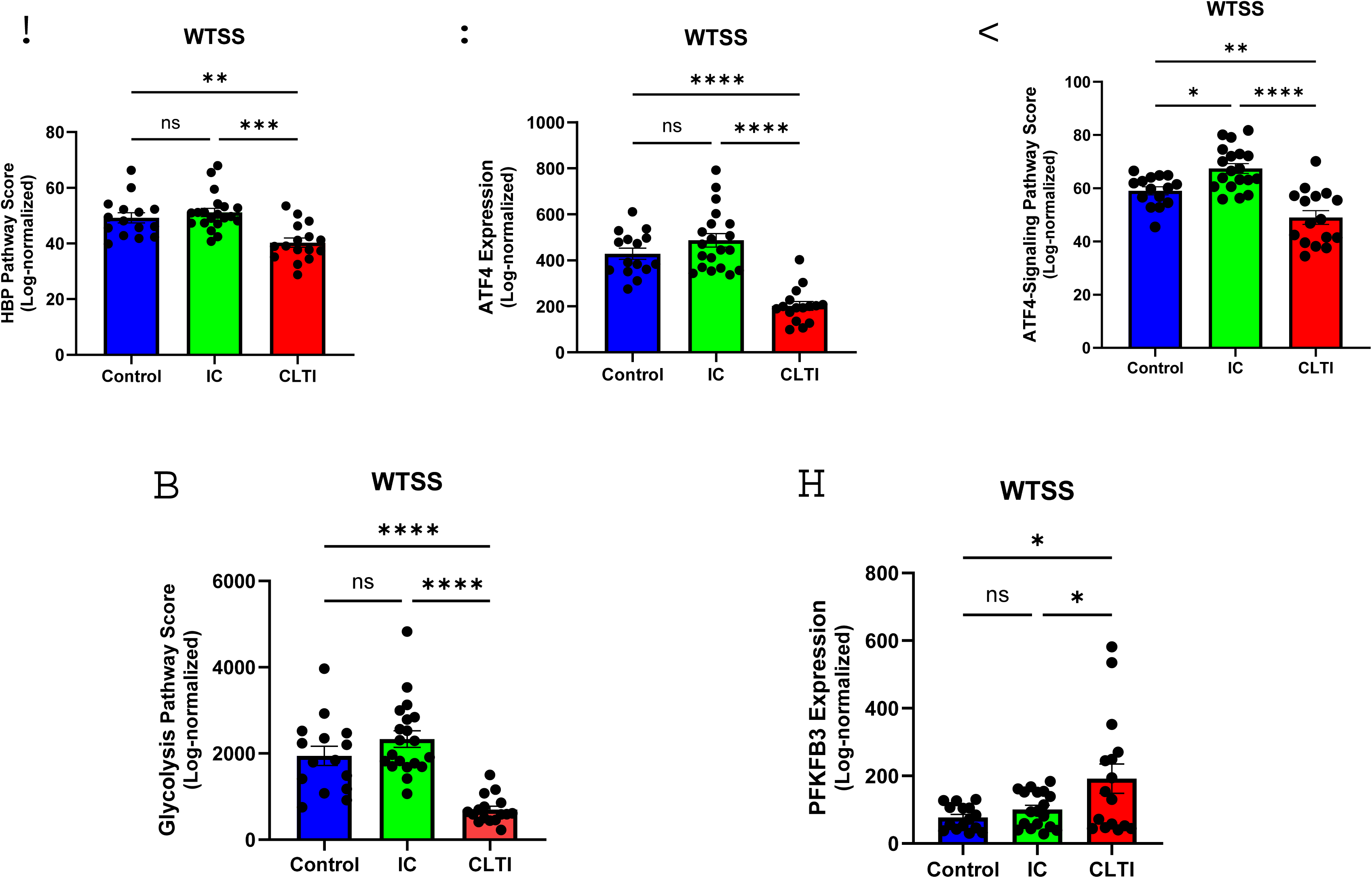
Bulk RNA sequencing of human lower limb muscle reveals metabolic reprogramming in the HBP–ATF4 axis and glycolysis in CLTI versus control and IC. **(A–E)** Quantification of log-normalized expression from whole transcriptome shotgun sequencing (WTSS) bulk RNA-seq data from gastrocnemius muscle of non-PAD controls, intermittent claudication (IC), and chronic limb-threatening ischemia (CLTI) patients. Composite pathway scores were computed as the average log-normalized expression of genes from each pathway. **(A)** HBP pathway score,**(B)** ATF4 expression,**(C)** ATF4 signaling pathway score,**(D)** Glycolysis pathway score,**(E)** PFKFB3 expression. Each dot represents expression level in a single human subject. Bars represent mean ± SEM. Comparisons were performed using one-way ANOVA with Tukey’s multiple comparisons test for normally distributed variables and Kruskal-Wallis test with Dunn’s post hoc test for non-normally distributed variables. Significance: ns = not significant; * <0.05; ** <0.01; *** <0.001; **** <0.0001. Groups: Control (blue), IC (green), CLTI (red). See also Supplementary Figure S1-3 for gene lists used in pathway score generation.

Expression of ATF4, a transcription factor integral to the integrated stress response (ISR) and a key regulator of ischemia-induced “exercise-like” angiogenesis mediated by HBP activation in PAD, was also significantly decreased in CLTI muscle (202.00 ± 19.02, mean ± SEM, n = 16) compared with both IC (486.90 ± 29.20, mean ± SEM, n = 20; P < 0.001) and control (428.80 ± 24.55, mean ± SEM, n = 15; P < 0.001; Figure 1B).^18,19^ A composite gene expression signature of genes downstream of the ATF4 transcriptional pathway was likewise significantly downregulated in CLTI (48.96 ± 2.55, mean ± SEM, n = 16) compared with IC (67.42 ± 1.83, mean ± SEM, n = 20; P < 0.001) and control (59.01 ± 1.49, mean ± SEM, n = 15; P = 0.004; Figure 1C). This reduction was driven by downregulation of several genes, including (ACOT2, CARS, CRISPLD2, CTH, EIF4EBP1, GARS, GOT1, GPT2, GTPBP2, LMO4, NARS, PHF10, SERPINF1, SHMT2, SLC19A2, SLC1A4, SLC25A33, TAF15, and TRIB3). Dot plots showing expression patterns of the individual genes within the ATF4 transcriptional pathway are shown in Supplementary Figure S3.

A different pattern emerged from the study of glycolysis. The overall glycolysis gene expression signature was significantly decreased in CLTI muscle (696.50 ± 80.16, mean ± SEM, n = 16) compared with both IC (2334 ± 192.70, mean ± SEM, n = 20; P < 0.0001) and control (1944 ± 221.70, mean ± SEM, n = 15; P < 0.0001; Figure 1D). This reduction was driven by downregulation of several genes, including (HK1, HK2, ALDOA, GAPDH, GPI, LDHA, PFKM, PGAM1, PGK1, PKM, and TPI1). Dot plots showing expression patterns of the individual genes within the glycolysis pathway are shown in Supplementary Figure S4. Among the PFKFB isoforms, PFKFB3 gene expression was significantly increased in CLTI muscle (191.90 ± 43.41, mean ± SEM, n = 16) compared with both Control (76.86 ± 9.55, mean ± SEM, n = 15; P = 0.012) and IC (100.50 ± 12.47, mean ± SEM, n = 18; P = 0.042; Figure 1E), consistent with prior findings.^21^ In contrast, PFKFB1, PFKFB2, and PFKFB4 were significantly reduced in CLTI compared with IC and controls (Supplementary Figure S5), highlighting selective upregulation of PFKFB3 in ischemic muscle.

We next analyzed the integrated single-cell RNA-seq (scRNA-seq) data to determine cell-type–specific expression of metabolic transcripts. We first characterized the cellular composition of the integrated scRNA-seq dataset derived from human skeletal muscle samples from control, IC, and CLTI patients. After performing unsupervised clustering of the integrated dataset, we identified 12 distinct clusters based on transcriptional profiles. These clusters were then annotated using canonical marker gene expression to define different cell populations, including endothelial cells, fibro-adipogenic progenitors (FAPs), mural cells, myonuclei, muscle stem cells (MuSCs), and immune cell subsets including T cells, B cells, NK cells, NKT cells, granulocytes, mast cells, and monocytes/macrophages (Figure 2A, B). No additional clusters met the criteria for a discrete, well-supported cell identity beyond these 12 populations.

**Figure 2.**
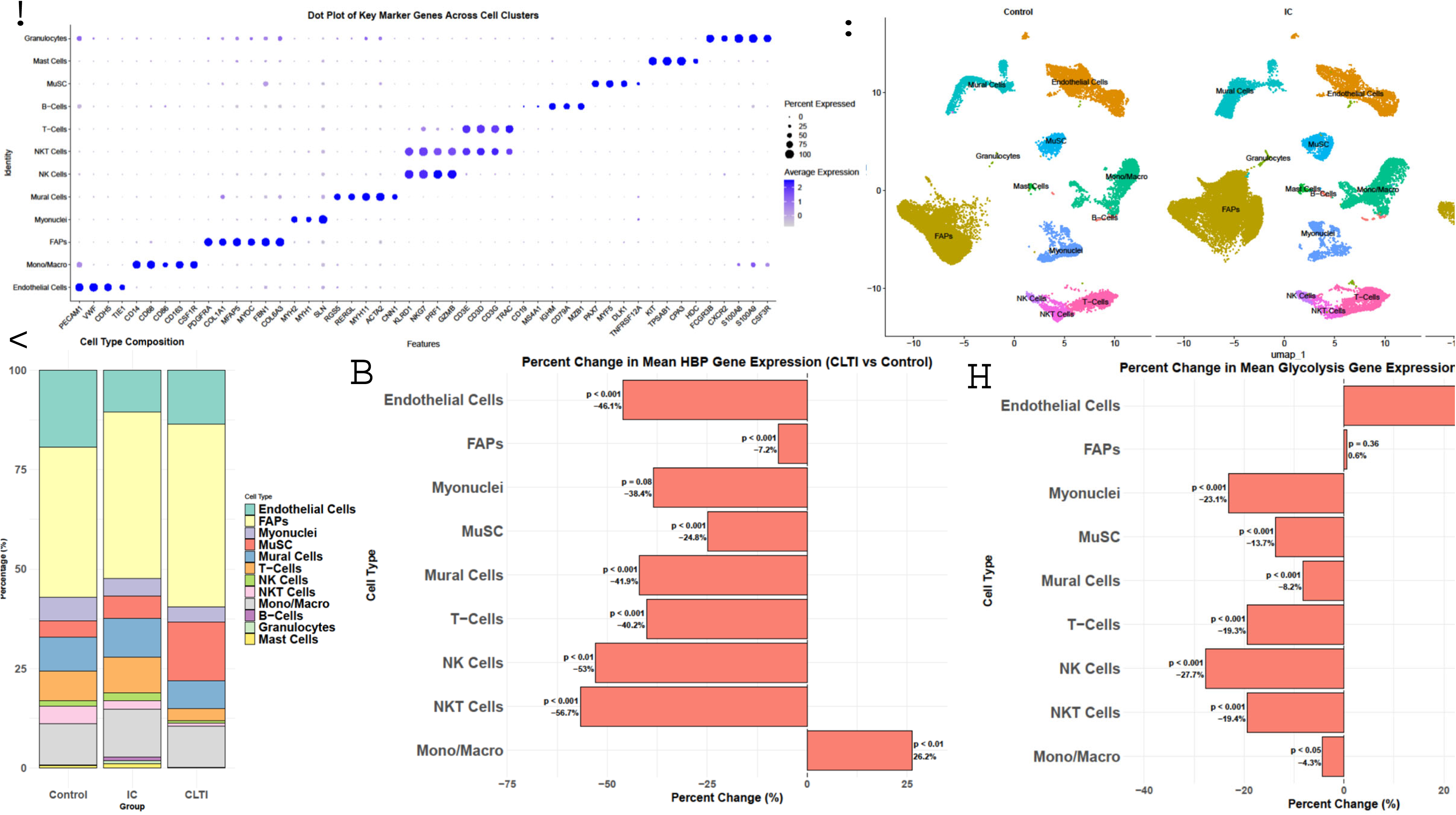
Integrated single-cell RNA sequencing analysis demonstrates cell type–specific alterations in metabolism in CLTI versus control. **(A)** Dot plot illustrating expression of key marker genes across identified clusters used to define cell type identities. Dot size represents the percentage of cells within each cluster expressing the gene; color intensity indicates average expression. **(B)** UMAP projection of Control, IC and CLTI muscle samples, showing distinct clusters including endothelial cells, fibro-adipogenic progenitors (FAPs), mural cells, myonuclei, monocytes/macrophages, and immune populations (T cells, B cells, NK cells, NKT cells, mast cells, granulocytes). **(C)** Cell type composition across groups (Control, IC, CLTI). Stacked bar plot shows the relative proportions of annotated cell populations per group. Each bar represents the average percentage of each annotated cell type (y-axis) within a group (x-axis). **(D)** Percent change in mean HBP gene expression (CLTI vs. Control) across cell types. **(E)** Percent change in mean glycolysis gene expression (CLTI vs. Control) across cell types. Percent change was computed as: ((Mean_CLTI – Mean_Control)/Mean_Control) × 100. Comparisons were performed using Wilcoxon rank-sum tests. Significance: *P < 0.05; **P < 0.01; ***P < 0.001; ns = not significant.

The frequency of skeletal muscle populations varied according to disease state (Figure 2C). FAPs constituted the largest compartment and progressively increased across groups (Control: 37.7%, IC: 41.8%, CLTI: 46.0%). Endothelial cells were reduced in both IC and CLTI relative to controls (19.3% in Control vs. 10.5% in IC and 13.6% in CLTI). Muscle cell myonuclei decreased in both IC and CLTI relative to controls (5.9% in Control, 4.5% in IC, 3.8% in CLTI), whereas MUSCs increased, particularly in CLTI (4.1% in Control, 5.6% in IC, 14.6% in CLTI). T cells displayed higher frequency in IC compared with Control but were reduced in CLTI (7.4% in Control, 8.9% in IC, 3.1% in CLTI). Cell types with representation below 1% in all groups (granulocytes, mast cells, B cells) were excluded from downstream metabolic analyses.

We next compared composite HBP and glycolysis gene expression scores across individual cell types. Pathway scores were calculated per cell and then averaged within each group (CLTI vs Control). In CLTI muscle, HBP gene expression was significantly reduced in endothelial cells (–46.1%, p < 0.001), mural cells (–41.9%, p < 0.001), T cells (–40.2%, p < 0.001), NK cells (–53.0%, p < 0.01), NKT cells (–56.7%, p < 0.001), muscle stem cells (–24.8%, p < 0.001), and FAPs (–7.2%, p < 0.001). Muscle cell myonuclei showed a non-significant decrease (–38.4%, p = 0.08). In contrast, monocytes/macrophages demonstrated a significantly increased HBP signature in CLTI vs control (+26.2%, p < 0.01) (Figure 2D). These findings indicate that HBP expression is significantly decreased in multiple structural and immune cell populations, while increased in monocytes/macrophages. This widespread reduction suggests that the global downregulation of HBP signaling observed in bulk tissue reflects downregulation across multiple cellular compartments.

Additionally, O-GlcNAc transferase (OGT) expression, the enzyme responsible for transferring UDP-GlcNAc to cytoplasmic and nuclear proteins, showed parallel patterns to HBP signature (Supplementary Figure S6A). Endothelial cells exhibited a significant reduction, with OGT levels decreased by 77% in CLTI compared with controls (fold change ∼0.23). Myonuclei, FAPs, and mural cells also showed lower expression in CLTI. In contrast, monocytes/macrophages demonstrated a significant increase in OGT expression in CLTI. These results indicate that OGT expression is reduced in different cell types, particularly endothelial cells, but increased in monocyte/macrophage in CLTI muscle vs control.

Glycolysis gene expression scores were also altered (Figure 2E). Endothelial cells showed a significant increased glycolysis in CLTI vs control (+36.3%, p < 0.001), consistent with previously reported findings in both cell culture and murine models of PAD.^16–18^ In contrast, glycolysis scores were significantly decreased in CLTI vs control in NK cells (–27.7%, p < 0.001), myonuclei (– 23.1%, p < 0.001), NKT cells (–19.4%, p < 0.001), T cells (–19.3%, p < 0.001), muscle stem cells (–13.7%, p < 0.001), mural cells (–8.2%, p < 0.001), and monocytes/macrophages (–4.3%, p < 0.05). However, FAPs did not significantly differ between groups (0.6%, p = 0.36). These results indicate that glycolysis gene expression is significantly increased in endothelial cells but decreased in most other structural and immune cell populations in CLTI muscle.

Regarding PFKFB3, data from the scRNA-seq was quite informative. We checked PFKFB3 expression across individual cell populations. In muscle cell myonuclei, PFKFB3 was reduced by approximately 32% (fold change ∼0.68), consistent with the decrease in glycolysis signature within the myonuclei. In contrast, PFKFB3 expression was increased across all other cell types, including endothelial cells. In endothelial cells, PFKFB3 expression was increased by approximately 225% (fold change ∼3.25), paralleling the elevated glycolysis gene signature observed in ECs (Supplementary Figure S6B).

Finally, to confirm that the single-cell data aligned with bulk observations, we compared group-level pathway scores averaged across all cells combined. Dot plots showing expression patterns for the genes in HBP and glycolysis pathways are shown in Supplementary Figures S7 and S8, respectively. Although myocytes were underrepresented in the single-cell dataset relative to whole-tissue RNA contributing to bulk data, group-level comparisons showed reduction in HBP and glycolysis signatures in CLTI compared to controls, consistent with the bulk RNA-seq results (Supplementary Figures S9A, B). These findings validate the bulk RNA-seq observations within the scRNA-seq dataset.

Together, these results reveal that metabolic alteration in PAD is not uniform across cell types. While global HBP and glycolysis signatures are broadly suppressed in CLTI muscle vs control, endothelial cells demonstrate increased glycolytic signature despite a parallel reduction in HBP signature, highlighting a potentially maladaptive metabolic phenotype in ischemic endothelium. A finding that could not be gleaned from analysis of the bulk RNA-seq data.

### Flow-Sorted Endothelial Cells from Ischemic Muscle in Patients with CLTI Exhibit Reduced OGT and Elevated PFKFB3 mRNA Expression Compared to Non-Ischemic Controls

To validate findings from our single-cell transcriptomic findings, we performed flow sorting of live endothelial cell populations from fresh muscle biopsies. Single-cell suspensions were gated to isolate CD56^-^CD31^+^ live ECs from CLTI muscle samples and non-PAD controls (Figure 3A). Patient demographics are provided in the Supplementary tables. The purity of the isolated endothelial fractions was confirmed by Western blotting for CD31 antibody, showing significant CD31 expression in sorted ECs comparable to HUVECs (used as a positive control), with absence of signal in myogenic progenitor cell lysates (negative control) (Figure 3B).

**Figure 3.**
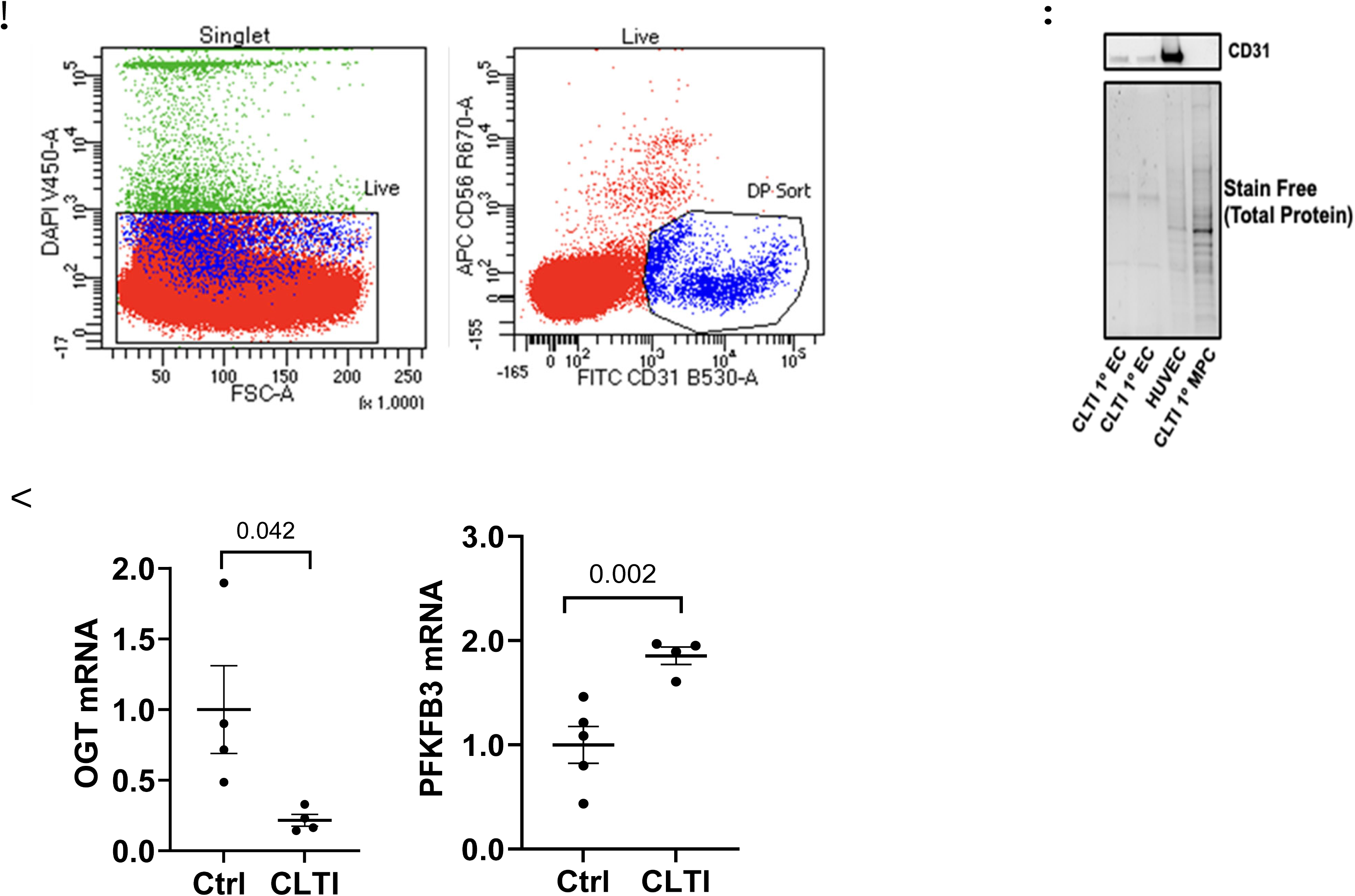
Flow-sorted endothelial cells from ischemic muscle confirm reduced OGT and elevated PFKFB3 expression in CLTI vs. controls. (A) Representative gating strategy for isolation of live CD56⁻CD31⁺ endothelial cells (ECs) from gastrocnemius muscle biopsies of CLTI patients and non-PAD controls. (B) Validation of flow-sorted primary ECs by immunoblot analysis for CD31. Human umbilical vein endothelial cells (HUVECs) served as a positive control, while myocyte progenitor cells (MPCs) served as a negative control. (C) Quantitative PCR analysis of sorted ECs from controls (n = 5) and CLTI (n = 4) patients showing relative expression of PFKFB3 and O-GlcNAc transferase (OGT). Statistical comparisons were performed using two-sample t-tests (unequal variances for OGT). Bars represent mean ± SEM. *P < 0.05; **P < 0.01; ns = not significant.

Quantitative PCR analysis of the sorted EC populations demonstrated a significant reduction in OGT mRNA levels in endothelial cells derived from CLTI muscle compared to controls (0.21 ± 0.04 vs. 1.00 ± 0.31, mean ± SEM, n= 4-5/group, p= 0.042, Figure 3C). In contrast, PFKFB3 mRNA levels were significantly higher in the ECs from patients with CLTI compared to controls (1.855 ± 0.08 vs. 1.00 ± 0.17, mean ± SEM, n= 4-5/group, p= 0.002, Figure 3C). These results validate the finding of the generated scRNA-seq data set.

### Immunofluorescence Validates Reduced O-GlcNAcylation and Elevated PFKFB3 Protein Expression in Endothelial Cells from Ischemic Muscle in Patients with CLTI Compared to Non-Ischemic Controls

We next sought to determine whether these transcriptional changes translated to protein level alterations. Therefore, we performed immunofluorescence staining to investigate the O-GlcNAcylation and PFKFB3 in endothelial cells of gastrocnemius (GA) muscle sections from 5 and 7 patients with CLTI vs 5 and 7 non-PAD controls respectively. Patient demographics are provided in the Supplementary tables. Antibodies against O-GlcNAC, CD31 (endothelial cell marker), and PFKFB3 were used as previously described.^18^ To exclude non-specific binding, secondary antibodies alone were used as negative controls on one section of each slide. A positive signal was defined by the presence of fluorescence in sections incubated with both primary and corresponding secondary antibodies but absent in sections incubated with secondary antibody alone. Any signal above the baseline threshold was considered a positive cell. Instances of multiple cells appearing clumped together were separated and counted, with signal intensity used to distinguish individual cells (Supplemental Figure S10). All imaging and quantification were performed blinded to group assignments to ensure unbiased analysis.

O-GlcNAC expression in ECs from tissue sections was quantified by counting cells co-stained with O-GlcNAC and CD31 in merged images, using the average from at least five random high-power fields (20× magnification) and expressed as the average number of O-GlcNAC/CD31 double-positive cells relative to the total CD31 within an individual patient. Similarly, PFKFB3⁺ ECs were quantified. To confirm CD31 specificity as an EC marker and rule out potential overlap with other CD31 expressing cell types, we performed co-staining of CD31 with CD45, a pan immune cell marker.^24^ Analysis revealed minimal overlap, with only 3.50 ± 1.08% of total CD31 staining being non-endothelial (mean ± SEM, n = 3 controls, Supplemental Figure S11 A). Macrophages may also express CD31. Therefore, we performed co-staining of CD31 with CD68, a macrophage marker. Similarly, this analysis showed minimal overlap, with only 1.48 ± 0.41% of total CD31 staining being non-endothelial (mean ± SEM, n = 3 controls, Supplemental Figure S11 B). These results collectively demonstrate that CD31 had very little overlap in expression with macrophage or immune cells.

Representative images showed reduced O-GlcNAc staining in CD31⁺ ECs from CLTI muscle (Figure 4A). Quantification confirmed a significant reduction in O-GlcNAc⁺ ECs in CLTI vs. controls (9.49 ± 3.33% vs. 31.33 ± 4.54%, mean ± SEM, n = 5/group, p= 0.004, Figure 4B). To determine whether this decrease in O-GlcNAC^+^ cells was merely due to differences in EC density, we quantified the total CD31 signal/area of tissue. We observed a numerically higher, but not statistically significant difference in capillary density/field area in patients with CLTI compared to controls (139.30 ± 21.88 vs. 108.70 ± 15.48, mean ± SEM, n = 5/group, p = ns, Figure 4C).

**Fig. 4.**
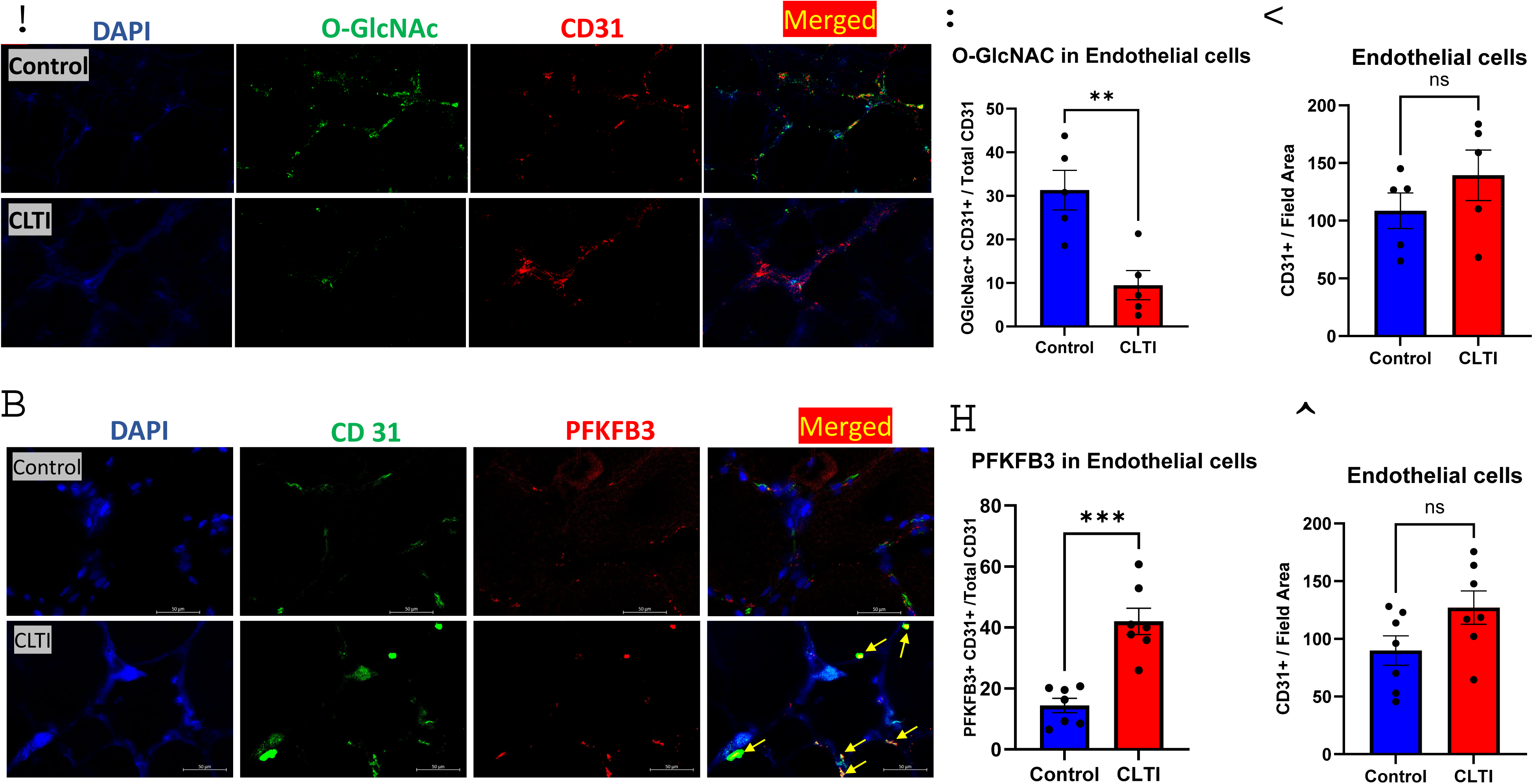
In calf skeletal muscle from patients with CLTI vs. controls CD-31+ cells show reduced O-GlcNAcylation but increased PFKFB3 expression. (**A**) DAPI (Blue), O-GlcNAC (Green), and CD31 (EC marker) (Red) immunostaining was performed on gastrocnemius (GA) muscle sections from PAD patients with chronic limb threatening ischemia (CLTI) and matched subjects without PAD (Control) (scale bar: 50 *μ*m). Representative images were taken with 40X magnification. (**B**) Quantitative analysis of colocalized O-GlcNAC with CD31 over total CD31 detected by immunofluorescence, (n=5/group). Unpaired t-test. (**C**) Quantitative analysis of capillary density (CD31 numbers) per magnification field (per 20X). detected by immunofluorescence, (n=5/group). Unpaired t-test. (**D**) DAPI (Blue), CD31 (EC marker) (Green), and PFKFB3 (Red) immunostaining was performed on GA muscle sections from PAD patients with CLTI and matched subjects without PAD (Control) (scale bar: 50 *μ*m). Representative images were taken with 40X magnification. (**E**) Quantitative analysis of colocalized PFKFB3 with CD31 over total CD31 detected by immunofluorescence, (n=7/group). Unpaired t-test. (**F**) Quantitative analysis of capillary density (CD31 numbers) per magnification field (per 20X). detected by immunofluorescence, (n=7/group). Unpaired t-test Data are presented as mean ± SEM, with individual data points shown. Statistical analyses were performed using unpaired t-tests. Significance: ns = not significant; * <0.05; ** <0.01; *** <0.001; **** <0.0001.

Representative images from CLTI muscle showed a greater PFKFB3 staining in CD31⁺ ECs (Figure 4D). Quantification confirmed PFKFB3⁺ ECs were significantly increased in CLTI muscle (42.01 ± 4.32% vs. 14.40 ± 2.32%, mean ± SEM, n = 7/group, p < 0.001, Figure 4E). Capillary density also showed a non-significant increase (127 ± 14.51 vs. 89.87 ± 12.79, mean ± SEM, n = 7/group, p = ns, Figure 4F).

These findings demonstrate that endothelial cells in ischemic muscle from CLTI patients exhibit both reduced O-GlcNAcylation and elevated PFKFB3 protein expression, confirming protein expression predicted from transcriptomic shifts.

### Exercise-induced angiogenesis in leg muscle in patients with IC is associated with decreased PFKFB3 in endothelial cells

In patients with PAD that can tolerate activity, supervised exercise is the only non-surgical intervention proven to enhance functional performance, with angiogenesis in the ischemic muscle occurring before clinical improvements are observed.^25,26^ In our recent report, we showed that O-GlcNAc-positive ECs were significantly increased in patients with IC following after 12 weeks of either supervised exercise plus Optimal Medical Care (OMC) vs. OMC alone.^18^ Therefore, we aimed to investigate the impact of supervised exercise on EC PFKFB3 in patients with IC. The study included two demographically comparable groups: patients with IC assigned to either exercise plus OMC (n=11) or OMC (n=9). Patient demographics are provided in the Supplementary tables. GA muscle sections from both groups were stained with antibodies against PFKFB3 and CD31 to evaluate ECs before and after 12 weeks of either supervised exercise plus OMC vs. OMC alone (Fig. 6A).

As previously reported, supervised exercise increases capillary density in ischemic GA muscles of IC patients compared to those receiving OMC alone.^25^ Consistent with our previous results, after 12 weeks, capillary density was significantly higher in the supervised exercise group (169.6 ± 14.32 vs. 113.3 ± 7.17, capillary density per magnification field, mean ± SEM, n=11/group, p=0.001), whereas no significant change was observed in the OMC group (120.3 ± 7.78 vs. 118.3 ± 6.83, mean ± SEM, n=9/group, p=0.57, Fig. 5A-B).^18,25,27^ We then assessed changes in PFKFB3 expression in ECs in response to supervised exercise versus OMC. At baseline, there was no significant difference between the exercise and OMC groups in the percentage of ECs (CD31^+^ cells) expressing PFKFB3 (14.62% ± 1.38 vs. 11.57% ± 1.65, mean ± SEM, n=11 for Exercise, n=9 for OMC, p=0.17, Fig. 6C). However, after 12 weeks, the percentage of PFKFB3-positive ECs remained unchanged in the OMC group (11.24% ± 1.68 vs. 11.57% ± 1.65, mean ± SEM, n=9, p=0.86, Fig. 6C). In contrast, supervised exercise led to a significant reduction in PFKFB3-positive ECs (2.67% ± 0.39 vs. 14.62% ± 1.38, mean ± SEM, n=11, p<0.001, Fig. 6C).

**Fig. 5.**
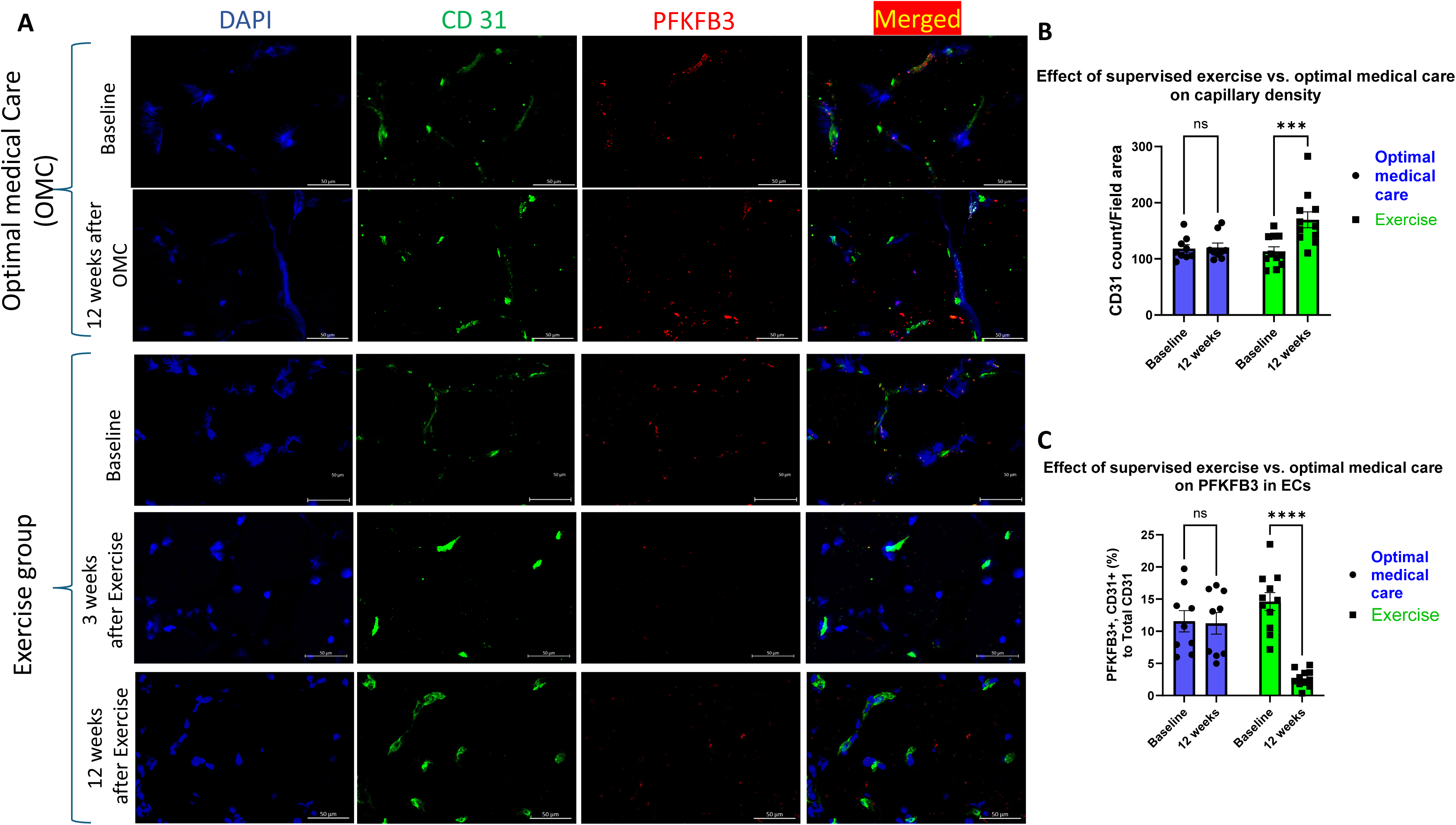
In patients with intermittent claudication, 12 weeks of supervised exercise plus optimal medical care (OMC) vs. OMC alone decreased PFKFB3 in endothelial cells. **(A)** Representative images of gastrocnemius muscle sections from patients with IC before and after 6 and 12 weeks of supervised exercise plus optimal medical care (OMC) (n=11) or OMC alone (n=9). Sections stained for DAPI (blue), CD31 (green), and PFKFB3 (red). Scale bar = 50 μm. **(B)** Quantification of capillary density (CD31 numbers) per magnification field (per 20X). **(C)** Quantification of colocalized PFKFB3 and CD31 to total CD31. Statistical comparisons between the groups in B and C were done using multiple comparisons Two-way ANOVA. Significance: ns = not significant; * <0.05; ** <0.01; *** <0.001; **** <0.0001.

Collectively, these findings demonstrate that supervised exercise in IC patients leads to increased capillary density while significantly reducing PFKFB3 expression in ECs. This suggests that exercise-induced vascular remodeling occurs through distinct metabolic pathways compared to those activated in CLTI, potentially explaining the beneficial effects of supervised exercise programs in PAD patients.

## Discussion

PAD is a major health care problem for which few medical therapies are able to alter outcomes for patients. Lower leg muscle is the “target” organ affected in PAD. To the best of our knowledge, this study presents the first scRNA-seq data that examines cell-specific expression from control subjects, patients with IC, and CLTI. This dataset was generated by combining two independent scRNA-seq datasets that were log-normalized and integrated using canonical correlation analysis and Harmony batch correction. By performing this analysis, the new dataset showed that ischemic ECs in subjects with CLTI undergo a maladaptive metabolic shift characterized by increased PFKFB3 and overall glycolytic gene expression, a conclusion that would not have been possible using bulk RNA sequencing. Of equal or greater importance, we were able to validate the hypotheses generated by cell sorting of ECs from a cohort of control and CLTI patients, as well as immunofluorescence staining of human muscle sections. Together, these data provide novel insights on the metabolic alterations found specifically in the ECs of CLTI patients’ limb tissues.

Though numerous studies can be found on the role of metabolism in controlling endothelial tumor angiogenesis, studies on EC metabolism in PAD are limited in number and only recently available.^3,16–18,28^ For example, the angiogenic response to HLI differs between in-bred mouse strains. C57BL/6J mice, which display a robust angiogenic response to HLI, showed no significant increase in EC glycolysis between the ischemic and non-ischemic limbs.^16,29^ In contrast, BALB/cJ mice, which exhibit a poor angiogenic response, generated a significant increase in glycolysis in ischemic vs. non-ischemic ECs.^16,29^ Treatment of BALB/cJ mice post-HLI with microRNA-93 improved perfusion recovery and normalized EC glycolytic flux by targeting increases in PFKFB3, whereas VEGF₁₆₅a treatment failed to improve perfusion despite increasing EC glycolysis.^16^ These data suggest that upregulated ischemia-induced glycolysis may be maladaptive, and that limiting increases in glycolytic flux may be a valuable therapeutic goal.

In ECs, VEGF₁₆₅a under hypoxia and serum starvation (HSS) induced PFKFB3 and glycolytic capacity, but also increased endothelial permeability and reactive oxygen species (ROS) generation.^16^ Conversely, microRNA-93 treatment of ECs under the same conditions suppressed greater PFKFB3 expression by targeting the mRNA. This attenuated increased glycolysis, and resulted in an upregulation of glucose-6-phosphate dehydrogenase (G6PD), thereby activating the PPP.^16^ Activating PPP under HSS led to increased production of NADPH, improved redox homeostasis, and reduced EC permeability under HSS. Finally, glucose bioavailability is not a limiting factor for perfusion recovery in ischemic muscle. In a murine model of PAD, intramuscular injection of D-glucose did not improve perfusion recovery compared to L-glucose (osmotic control) following HLI, suggesting that simply altering glucose availability is insufficient to drive therapeutic angiogenesis.^17^

In an alternative metabolism study, we recently reported that activation of the HBP pathway directly via glucosamine increased O-GlcNAcylation in ECs in-vitro, ECs from ischemic muscle, and in whole ischemic muscle. Increased O-GlcNAcylation increased EC growth without increasing permeability or ROS and this was mediated through ATF4; a transcription factor that has been shown to be linked to exercise-induced angiogenesis.^18,19^ Within the same report, HBP activation suppressed the otherwise expected increases in glycolysis, providing a hereunto unappreciated potential benefit.^18^ Both bulk and sc-RNA sequencing suggested reduced EC O-GlcNAcylation in patients with PAD (both IC and CLTI) vs. control, confirming that expression of genes across the HBP pathway was decreased, as was the key down-stream mediator of glucosamine, ATF-4.^18,19^ Additionally, supervised exercise, the only clinically approved non-surgical approach for improving functional performance in patients with PAD, has been shown to increase endothelial O-GlcNAcylation in human muscle.^18^ Though not from single data sequencing data, our studies from tissue sections showed that in patients with IC, supervised exercise vs. usual medical care suppressed the number of PFKFB3^+^ ECs.

In contrast to ECs, myonuclei in CLTI muscle exhibited downregulation of both PFKFB3 and glycolytic gene signatures, contributing to the overall suppression of glycolysis observed in bulk RNA-seq. This finding aligns with prior studies demonstrating that primary muscle cells derived from patients with CLTI exhibit reduced PFKFB3 expression and diminished glycolytic capacity, which were associated with compromised mitochondrial metabolism.^21^

Together, our paper used a newly created transcriptomic dataset and showed that alterations in metabolism in muscle of patients with PAD vs. control are not uniform across cell types. While global HBP and glycolysis signatures are broadly suppressed in CLTI muscle, ischemic endothelium from CLTI patients uniquely demonstrates a maladaptive metabolic shift characterized by suppressed O-GlcNAcylation and elevated glycolysis. These cell type–specific alterations would be expected to contribute to EC dysfunction in CLTI vs. control. Bidirectional targeting, suppressing EC glycolysis and enhancing muscle glycolysis, would be needed to improve outcomes in CLTI.

---

**Table 1:**
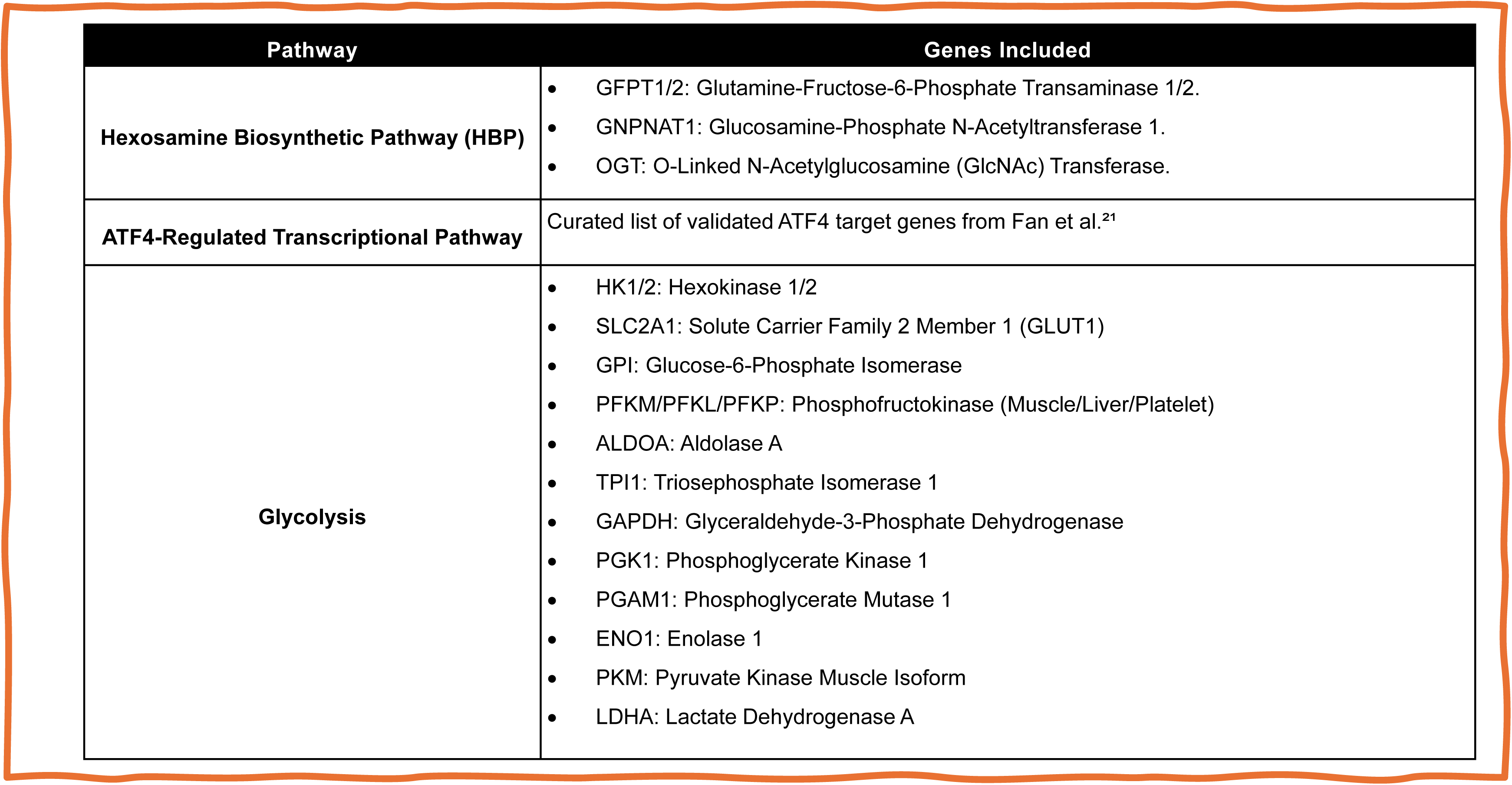
Genes Used to Generate Pathway Scores in Bulk and Single-Cell RNA-seq Analyses.

| Pathway | Genes Included |
| --- | --- |
| Hexosamine Biosynthetic Pathway (HBP) | GFPT1/2: Glutamine-Fructose-6-Phosphate Transaminase 1/2.<br>GNPNAT1: Glucosamine-Phosphate N-Acetyltransferase 1.<br>OGT: O-Linked N-Acetylglucosamine (GlcNAc) Transferase. |
| ATF4-Regulated Transcriptional Pathway | Curated list of validated ATF4 target genes from Fan et al.<sup>21</sup> |
| Glycolysis | HK1/2: Hexokinase 1/2<br>SLC2A1: Solute Carrier Family 2 Member 1 (GLUT1)<br>GPI: Glucose-6-Phosphate Isomerase<br>PFKM/PFKL/PFKP: Phosphofructokinase (Muscle/Liver/Platelet)<br>ALDOA: Aldolase A<br>TPI1: Triosephosphate Isomerase 1<br>GAPDH: Glyceraldehyde-3-Phosphate Dehydrogenase<br>PGK1: Phosphoglycerate Kinase 1<br>PGAM1: Phosphoglycerate Mutase 1<br>ENO1: Enolase 1<br>PKM: Pyruvate Kinase Muscle Isoform<br>LDHA: Lactate Dehydrogenase A |

**Table 2:** Canonical Marker Genes Used for Annotation of Major Cell Types in Human Skeletal Muscle Integrated scRNA-seq Dataset.

| Cell Type | Marker Genes |
| --- | --- |
| Endothelial cells | PECAM1, VWF, CDH5, TIE1, CLDN5 |
| Monocytes/Macrophages | CD14, CD68, CD86, CD163, CSF1R |
| Fibro-adipogenic progenitors (FAPs) | PDGFRA, COL1A1, MFAP5, MYOC, FBN1, COL6A3 |
| Myonuclei | MYH2, MYH1, SLN |
| Mural cells | RGS5, RERGL, MYH11, ACTA2, CNN1 |
| NK cells | KLRD1, NKG7, PRF1, GZMB |
| T cells | CD3E, CD3D, CD3G, TRAC |
| B cells | CD19, MS4A1, IGHM, CD79A, MZB1 |
| Muscle stem cells (MuSCs) | PAX7, MYF5, DLK1, TNFRSF12A |
| Mast cells | KIT, TPSAB1, CPA3, HDC |
| Granulocytes | FCGR3B, CXCR2, S100A8, S100A9, CSF3R |

**Table 3:** Demographics of CLTI and Control Patient Samples Used for Flow Cytometric Endothelial Cell Sorting.

|  | Controls | CLTI |
| --- | --- | --- |
| Number | 5 | 4 |
| Age (years) (mean ± SEM) | 55.4 ± 5.06 | 59.75 ± 4.71 |
| Race % Caucasians | 80% (4/5) | 75% (3/4) |
| Sex (Men/Women)<br>(as self reported) | 2/3 | 1/3 |
| Smoking history (previous) | -- | 75% (3/4) |
| Diabetes | -- | 75% (3/4) |
| Hypertension | -- | 75% (3/4) |
| Hyperlipidemia | -- | 75% (3/4) |
| Coronary Artery Disease | -- | 25% (1/4) |
| Chronic Kidney Disease | -- | 75% (3/4) |

**Table 4:** Demographics of CLTI and Control Patient Samples Used for CD31 and O-GlcNAc Immunofluorescence.

|  | Controls | CLTI |
| --- | --- | --- |
| Number | 5 | 5 |
| Age (years) (mean ± SEM) | 62.4 ± 3.43 | 68.6 ± 3.46 |
| Race % Caucasians | 100% (5/5) | 40% (2/5) |
| Sex (Men/Women)<br>(as self reported) | 3/2 | 2/3 |
| Smoking history (previous) | -- | 80% (4/5) |
| Diabetes | -- | 80% (4/5) |
| Hypertension | -- | 100% (5/5) |
| Hyperlipidemia | -- | 60% (3/5) |
| Coronary Artery Disease | -- | 20% (1/5) |
| Chronic Kidney Disease | -- | 80% (4/5) |

**Table 5:** Demographics of CLTI and Control Patient Samples Used for CD31 and PFKFB3 Immunofluorescence.

|  | Controls | CLTI |
| --- | --- | --- |
| Number | 7 | 7 |
| Age (years) (mean ± SEM) | 57 ± 2.58 | 65.43 ± 2.26 |
| Race % Caucasians | 42.8% (3/7) | 42.8% (3/7) |
| Sex (Men/Women)<br>(as self reported) | 3/4 | 5/2 |
| Smoking history (previous) | 28.5% (2/7) | 71.4% (5/7) |
| Diabetes | N/a | 71.4% (5/7) |

**Table 6:**
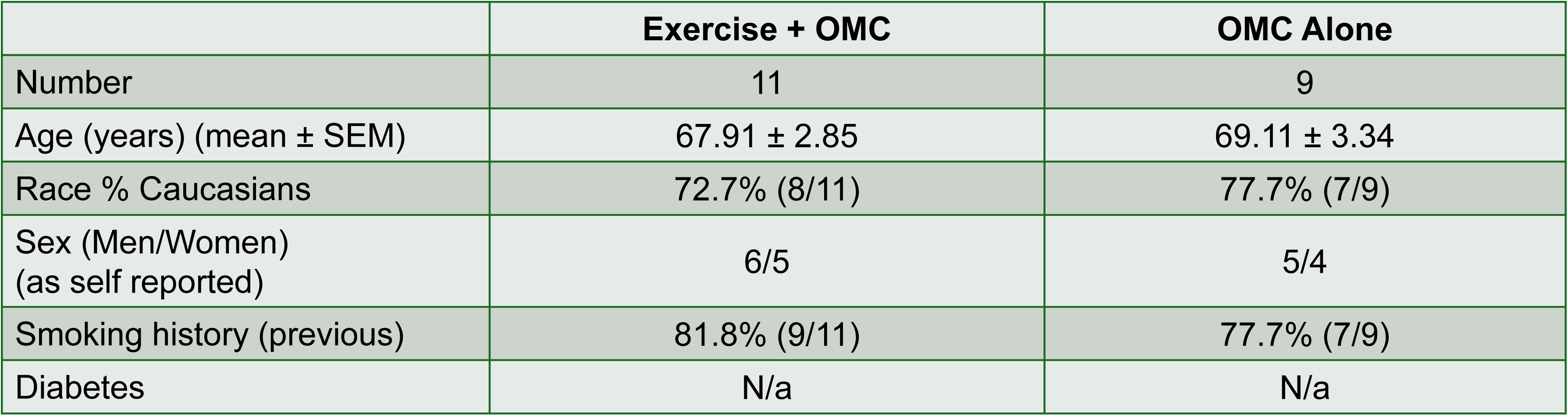
Demographics of Intermittent Claudication Patients in Supervised Exercise Versus Optimal Medical Care (OMC) Groups.

**Figure S1.**
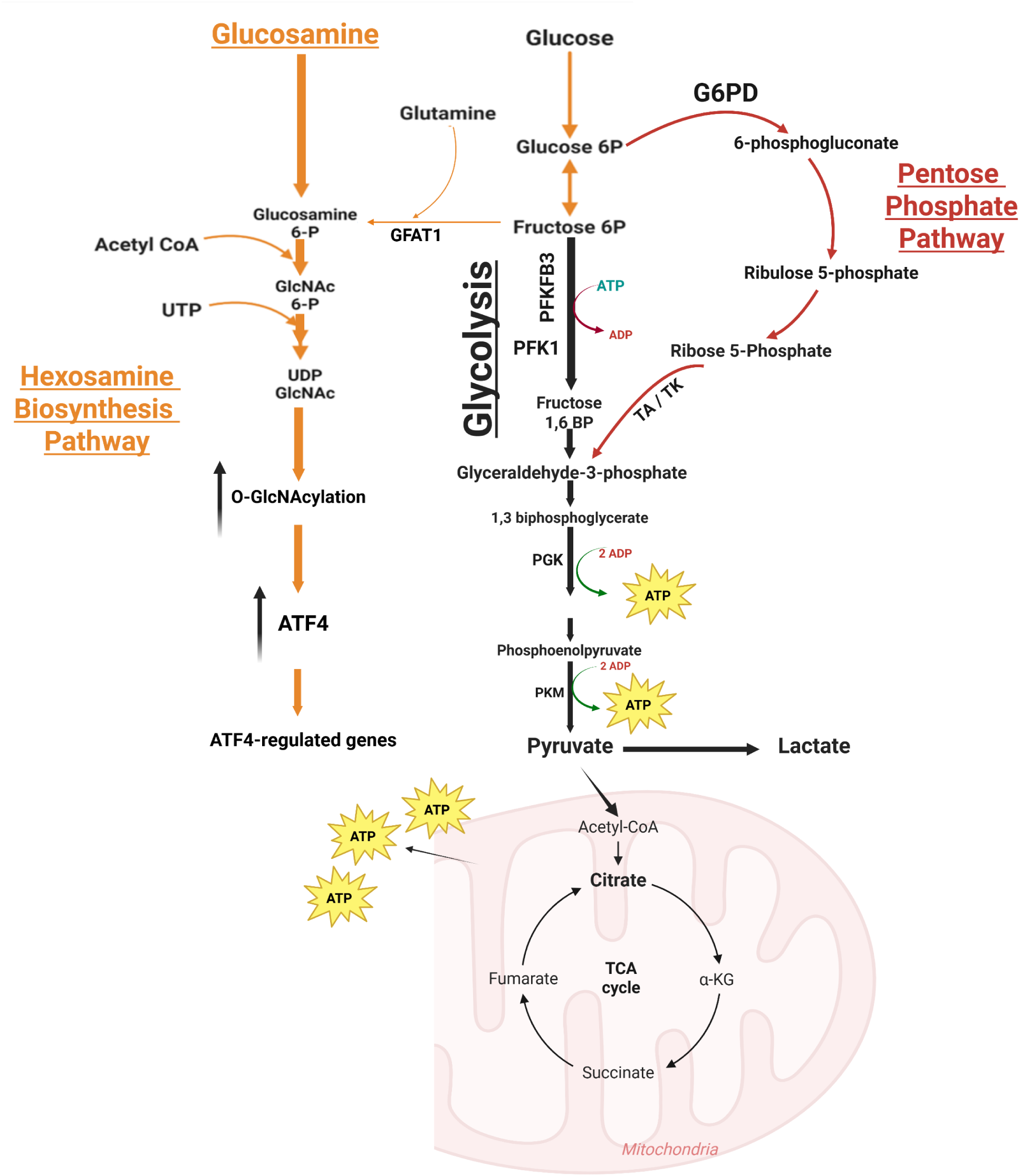
Schematic overview of glucose metabolism pathways in skeletal muscle and endothelial cells. This diagram illustrates the metabolic fate of glucose in skeletal muscle and endothelial cells, emphasizing three major pathways: Glycolysis (black pathway): Glucose is converted to glucose-6-phosphate (Glucose 6P), then to fructose-6-phosphate (Fructose 6P), and ultimately into pyruvate through the glycolytic cascade, generating ATP. The regulatory steps include PFK1 and PFKFB3, which promote the formation of fructose 1,6-bisphosphate (Fructose 1,6 BP). ATP generation is indicated at steps catalyzed by PGK and PKM enzymes. Pyruvate can be further oxidized in the mitochondrial TCA cycle (lower right) or converted to lactate. Hexosamine Biosynthesis Pathway (HBP; orange pathway): Fructose-6P is also a precursor for the HBP, where it is converted by GFAT1 into glucosamine-6P, ultimately forming UDP-GlcNAc, a key substrate for protein O-GlcNAcylation. This process can be activated by glucosamine supplementation, which bypasses GFAT1. Pentose Phosphate Pathway (PPP; red pathway): Alternatively, glucose-6P may enter the PPP through G6PD to generate ribose-5-phosphate for nucleotide biosynthesis and NADPH for redox balance. Abbreviations:

- PFK1: Phosphofructokinase 1, PFKFB3: 6-Phosphofructo-2-kinase/fructose-2,6-bisphosphatase 3, GFAT1: Glutamine:fructose-6-phosphate amidotransferase 1, G6PD: Glucose-6-phosphate dehydrogenase, PGK: Phosphoglycerate kinase, PKM: Pyruvate kinase M, TA: Transaldolase, TK: Transketolase.
- Glucose 6P: Glucose-6-phosphate, Fructose 6P: Fructose-6-phosphate, Fructose 1,6 BP: Fructose-1,6-bisphosphate, GlcNAc 6-P: N-acetylglucosamine-6-phosphate, UDP-GlcNAc: Uridine diphosphate N-acetylglucosamine, Ribose 5P: Ribose-5-phosphate, α-KG: Alpha-ketoglutarate, Acetyl-CoA: Acetyl-coenzyme A, ATP/ADP: Adenosine triphosphate/diphosphate.

**Figure S2.**
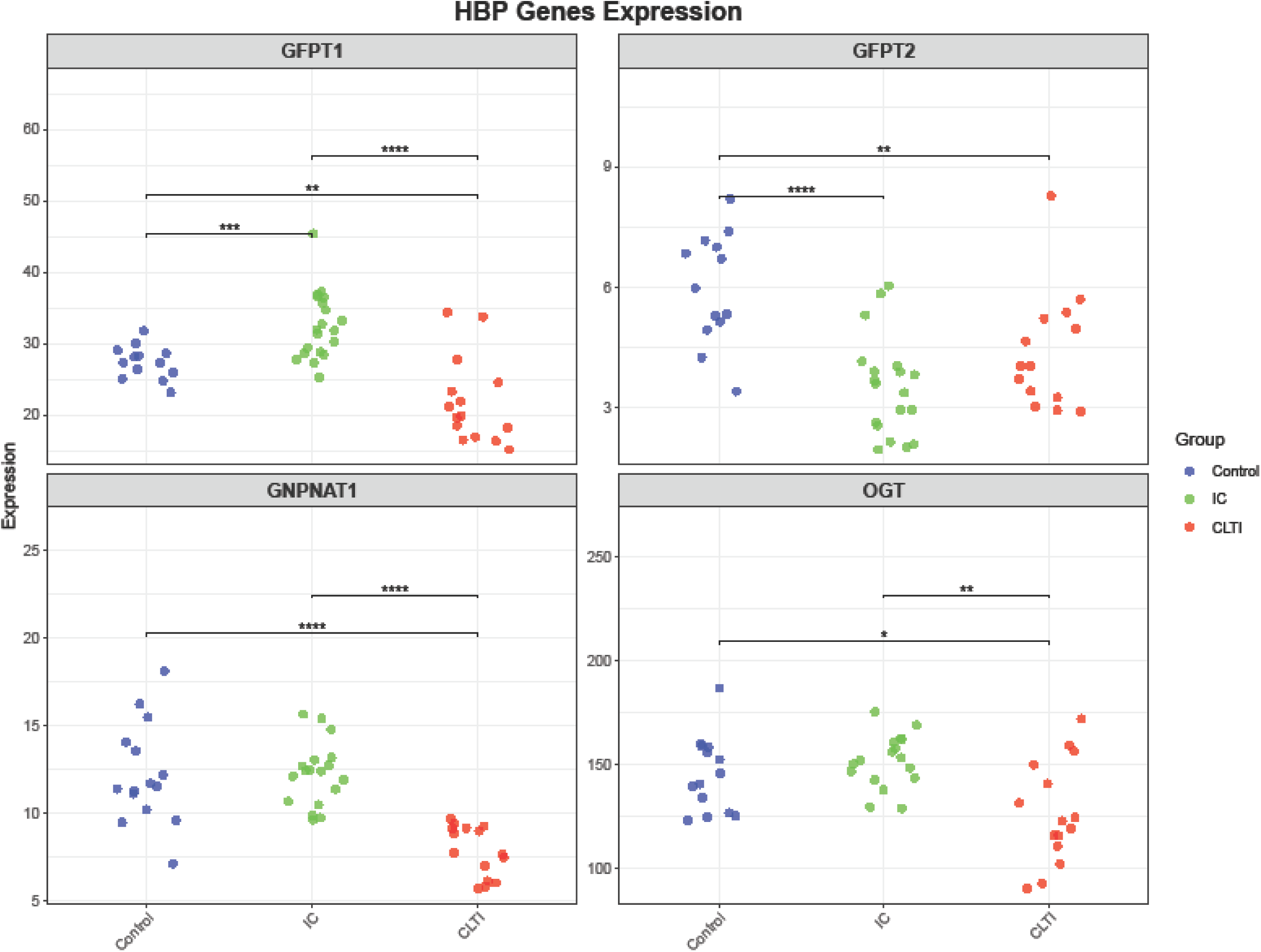
Expression of HBP genes in human skeletal muscle by bulk RNA sequencing. Dot plots show log-normalized expression of HBP genes (GFPT1, GFPT2, GNPNAT1, OGT) in gastrocnemius muscle samples from non-PAD controls (blue), intermittent claudication (IC, green), and chronic limb-threatening ischemia (CLTI, red). Each dot represents a single human subject. Genes are displayed in individual panels. Statistical analysis: one-way ANOVA with Tukey’s post hoc test. Significance: ns = not significant; *P < 0.05; **P < 0.01; ***P < 0.001; ****P < 0.0001.

**Figure S3.**
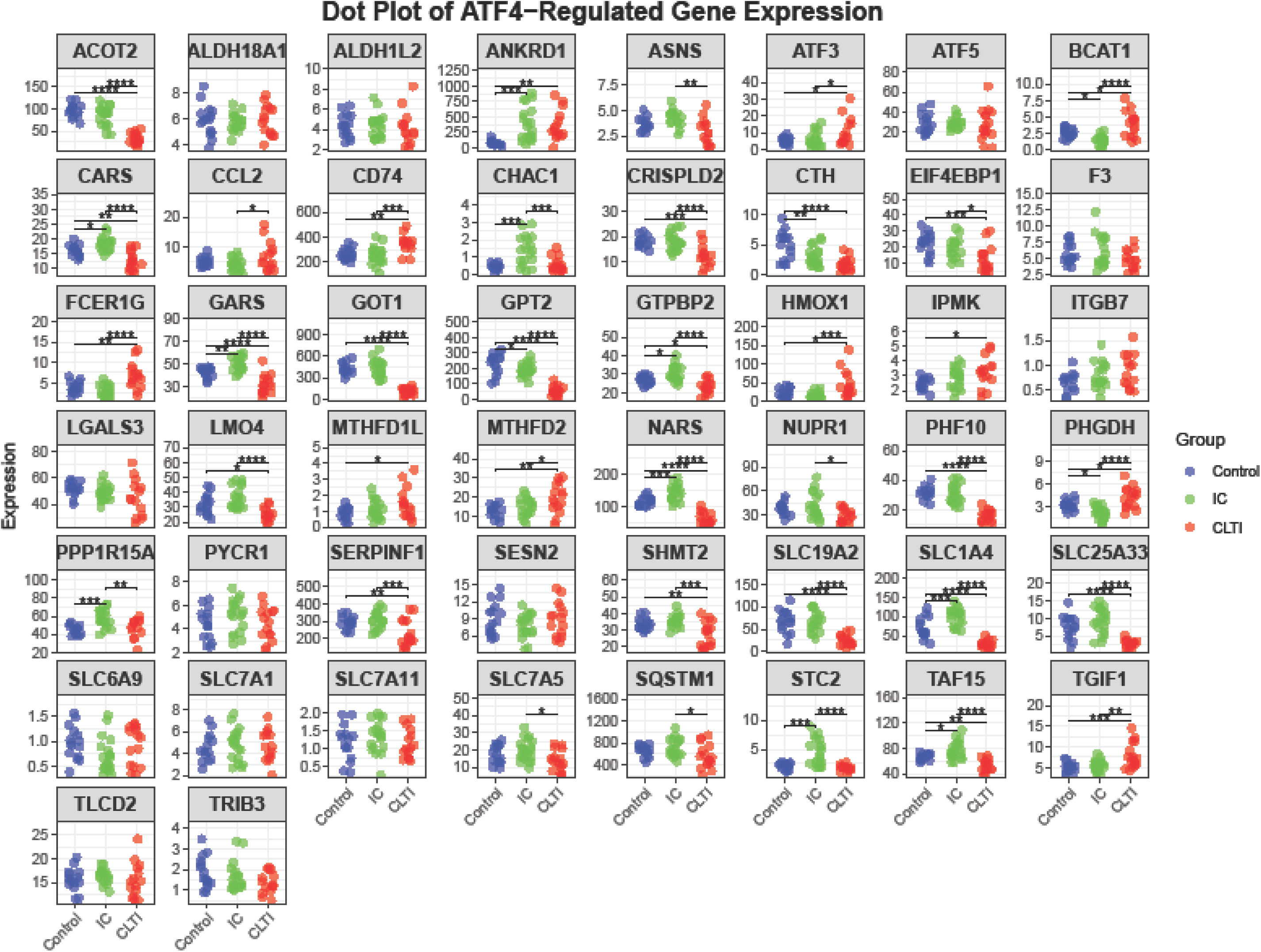
Expression of ATF4-regulated target genes in human skeletal muscle by bulk RNA sequencing. Dot plots show log-normalized expression of ATF4-regulated genes in gastrocnemius muscle samples from controls (blue), IC (green), and CLTI (red). Each dot represents a single human subject. Genes are displayed in individual panels. Statistical analysis: one-way ANOVA with Tukey’s post hoc test. Significance: ns = not significant; *P < 0.05; **P < 0.01; ***P < 0.001; ****P < 0.0001.

**Figure S4.**
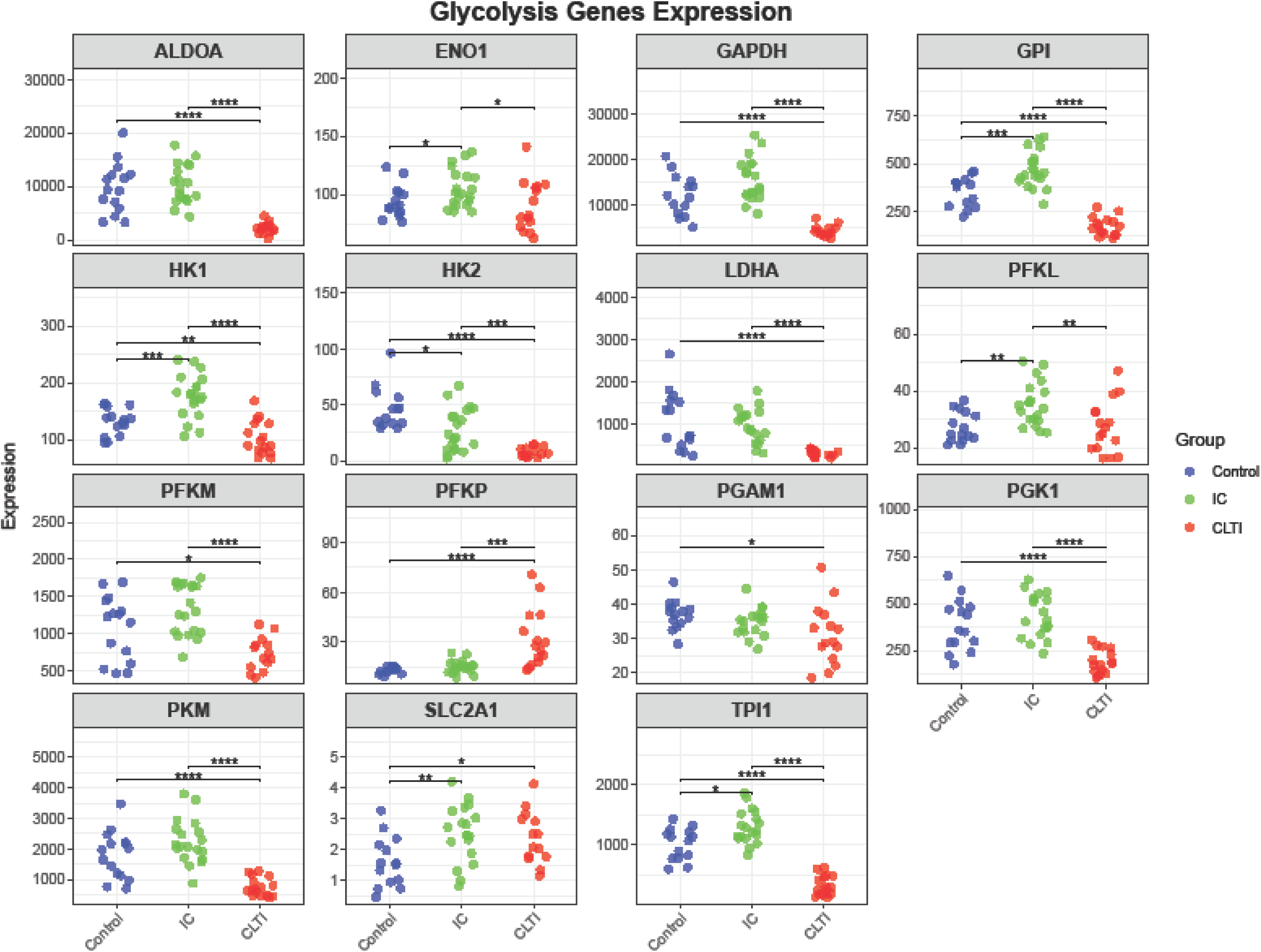
Expression of glycolytic genes in human skeletal muscle by bulk RNA sequencing. Dot plots of log-normalized expression of glycolytic pathway genes in gastrocnemius muscle from controls (blue), IC (green), and CLTI (red). Each dot represents a single human subject. Genes are displayed in individual panels. Statistical testing by one-way ANOVA with Tukey’s post hoc test. Significance: ns = not significant; *P < 0.05; **P < 0.01; ***P < 0.001; ****P < 0.0001.

**Figure S5.**
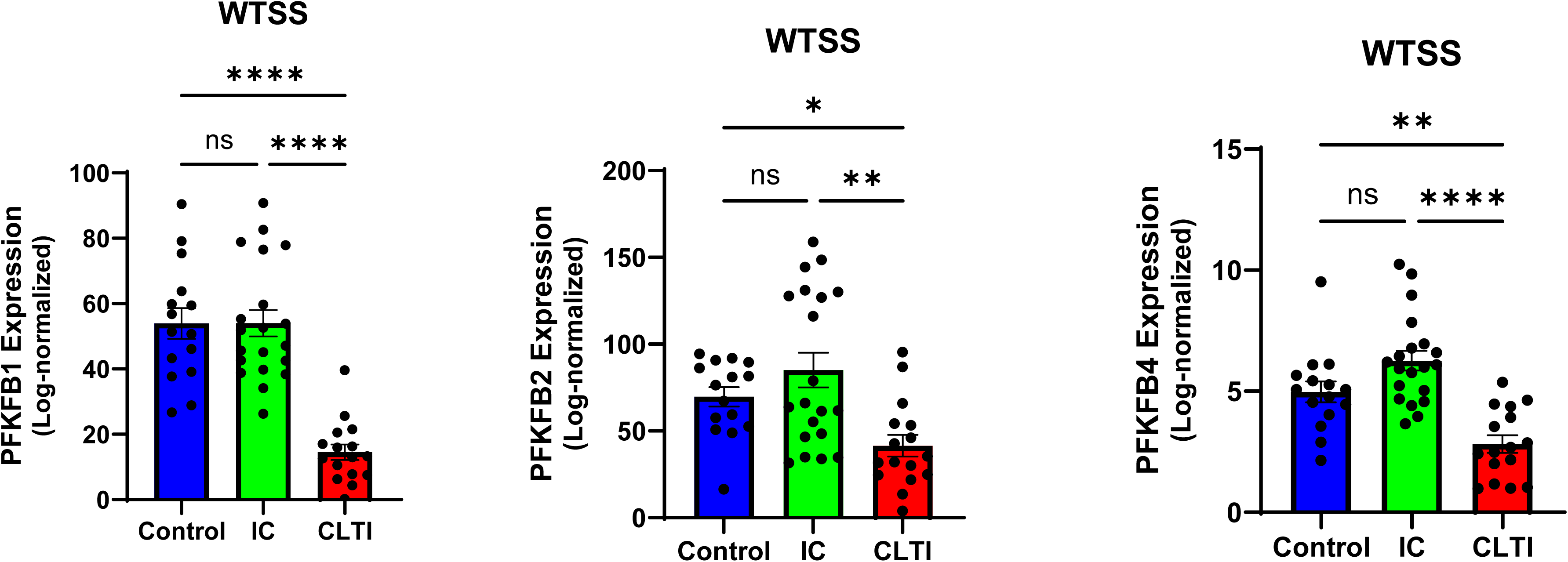
Expression of PFKFB isoforms in human skeletal muscle by bulk RNA sequencing. Dot plots of log-normalized expression of PFKFB1, PFKFB2, and PFKFB4 in gastrocnemius muscle from controls (blue), IC (green), and CLTI (red). Each dot represents a single human subject. Genes are displayed in individual panels. Statistical testing by one-way ANOVA with Tukey’s post hoc test. Significance: ns = not significant; *P < 0.05; **P < 0.01; ***P < 0.001; ****P < 0.0001.

**Figure S6.**
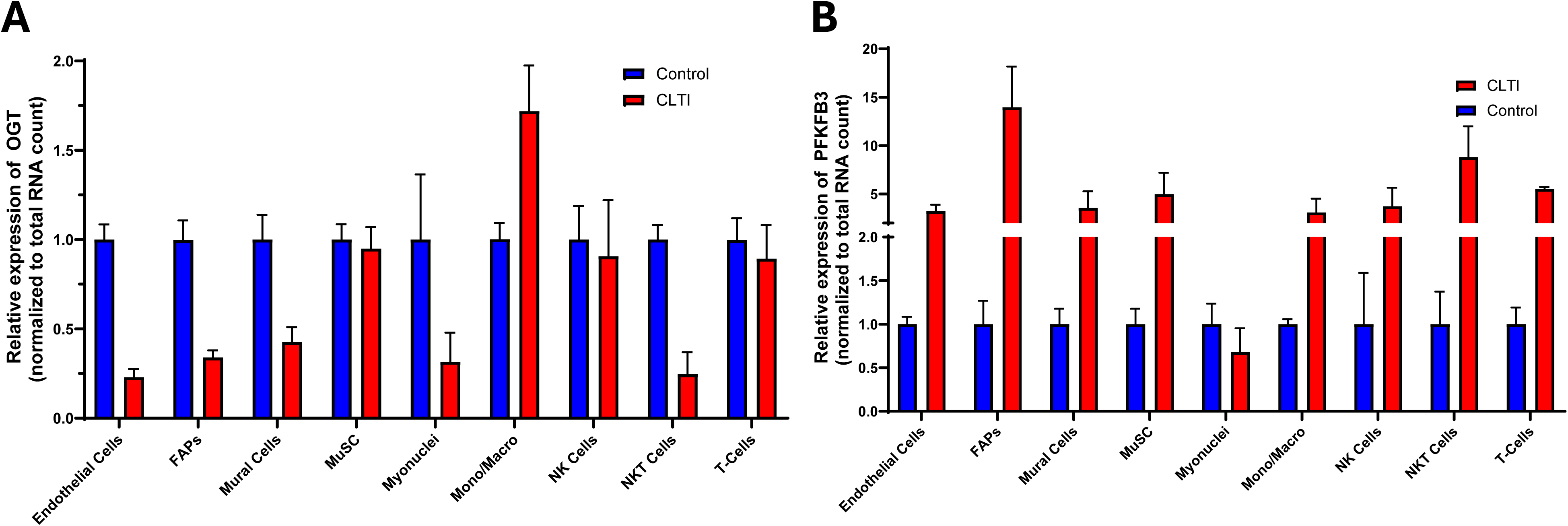
Single-cell RNA sequencing reveals reduced OGT and increased PFKFB3 in CLTI skeletal muscle vs controls. Integrated scRNA-seq of lower limb muscle shows (A) OGT expression and (B) PFKFB3 expression across major cell types in controls (blue) and CLTI (red). Data are mean ± SEM. Cell types include endothelial cells, fibro-adipogenic progenitors (FAPs), mural cells, muscle stem cells (MuSCs), myonuclei, monocytes/macrophages, B cells, NK cells, NKT cells, and T cells.

**Figure S7.**
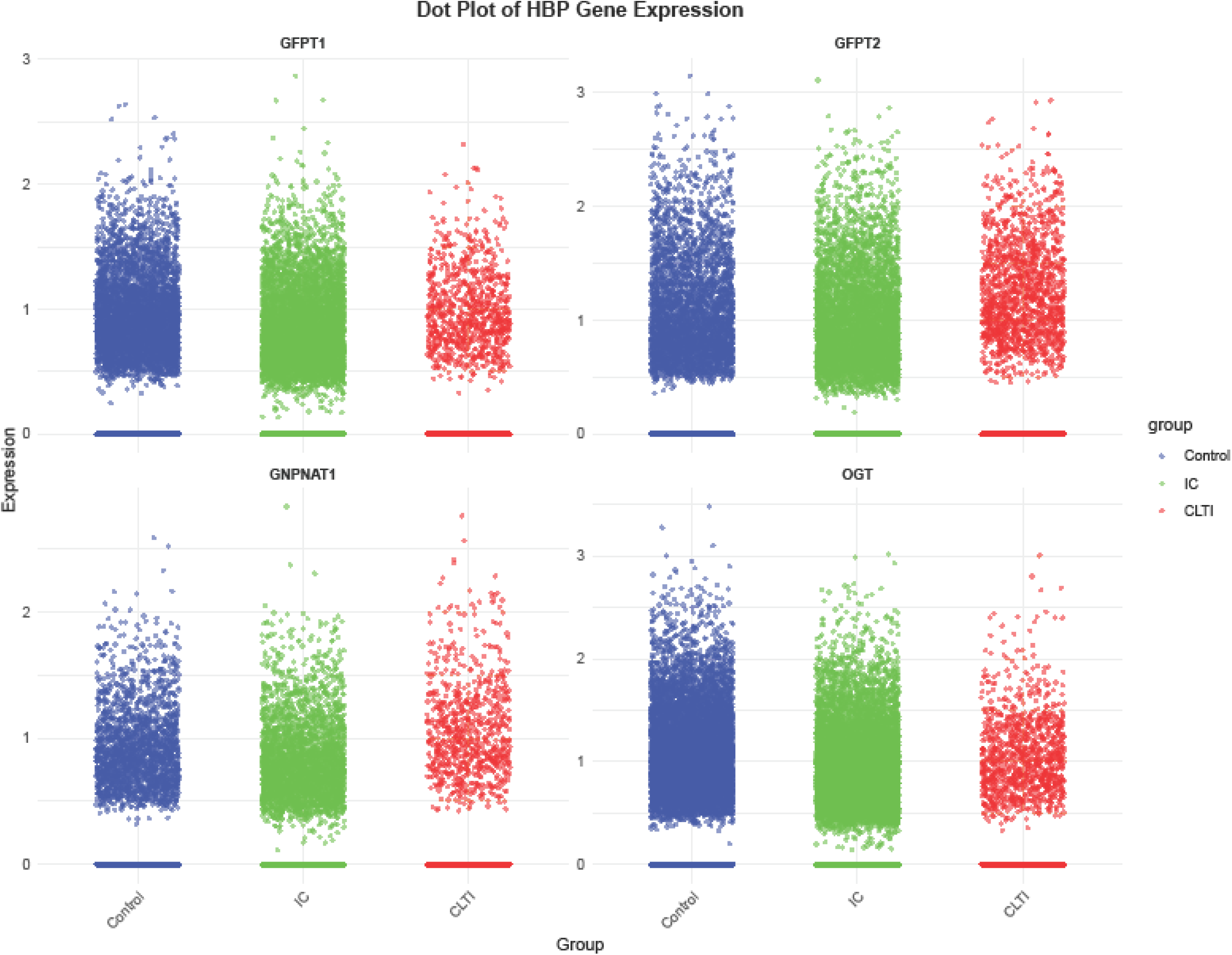
Expression of HBP genes in human skeletal muscle by the Integrated scRNA-seq. Dot plots of log-normalized expression of HBP genes (GFPT1, GFPT2, GNPNAT1, and OGT) from integrated scRNA-seq of lower limb muscle from controls (blue), IC (green), and CLTI (red). Each dot represents the expression value of a single cell.

**Figure S8.**
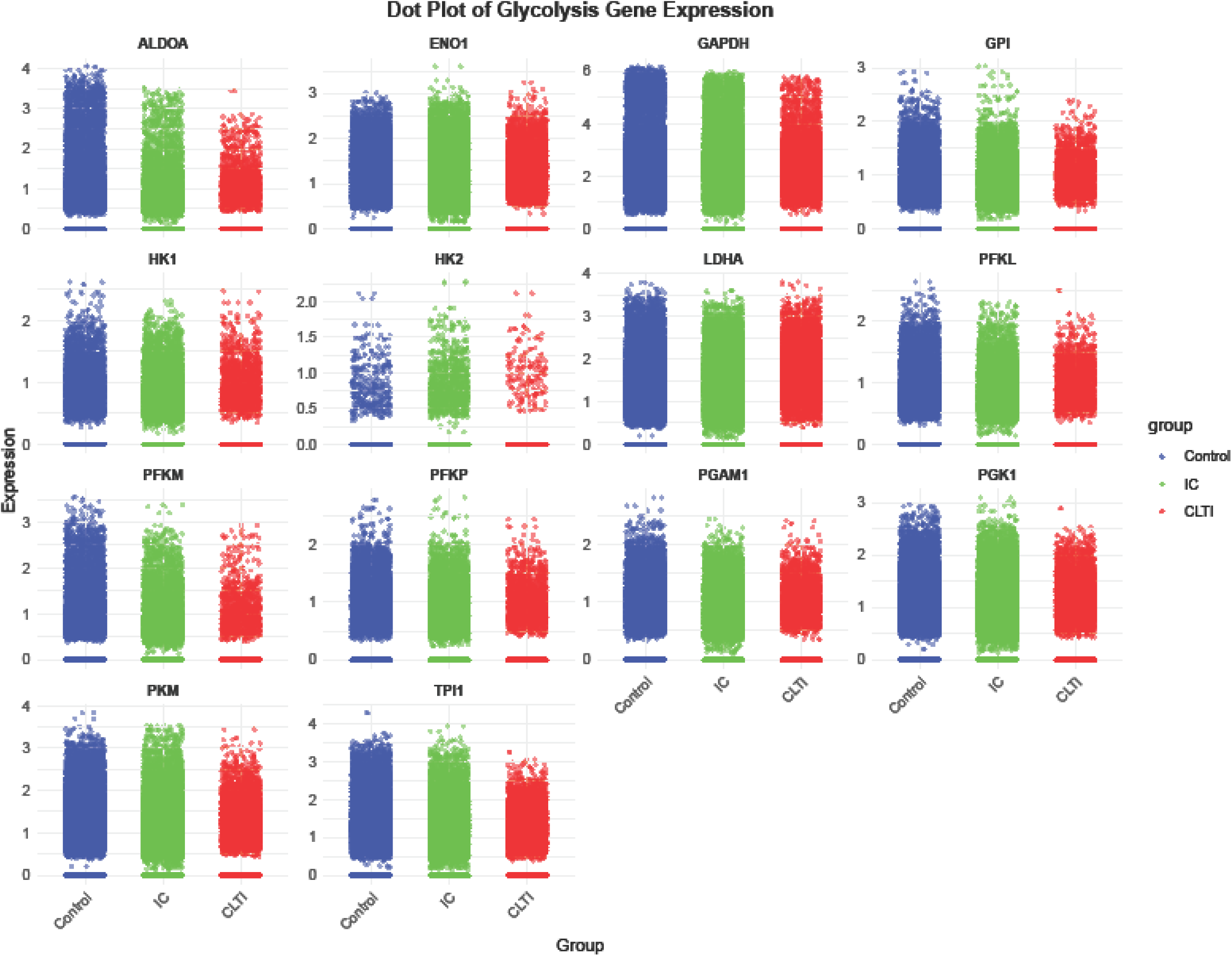
Expression of glycolytic genes in human skeletal muscle by the Integrated scRNA-seq. Dot plots of log-normalized expression of glycolysis-related genes (ALDOA, ENO1, GAPDH, GPI, HK1, HK2, LDHA, PFKL, PFKM, PFKP, PGAM1, PGK1, PKM, SLC2A1, TPI1) from integrated scRNA-seq of lower limb muscle from controls (blue), IC (green), and CLTI (red). Each dot represents the expression value of a single cell.

**Figure S9.**
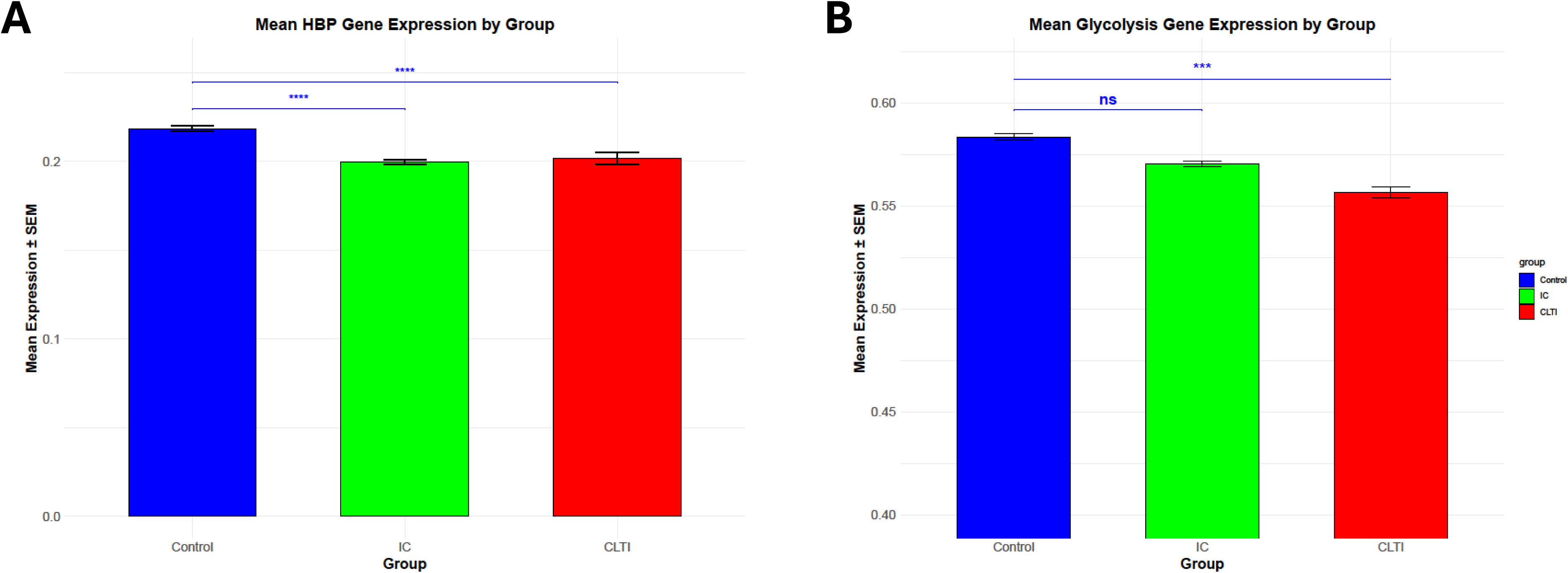
Pseudo-bulk analysis of integrated scRNA-seq data confirms bulk RNA-seq findings. **(A)** Bar plot of mean HBP gene expression across all cells in integrated scRNA-seq of lower limb muscle from controls (blue), IC (green), and CLTI (red). **(B)** Bar plot of mean glycolysis gene expression across the same dataset. Error bars represent the standard error of the mean (± SEM). Significance: ns = not significant; *P < 0.05; **P < 0.01; ***P < 0.001; ****P < 0.0001..

**Figure. S10.**
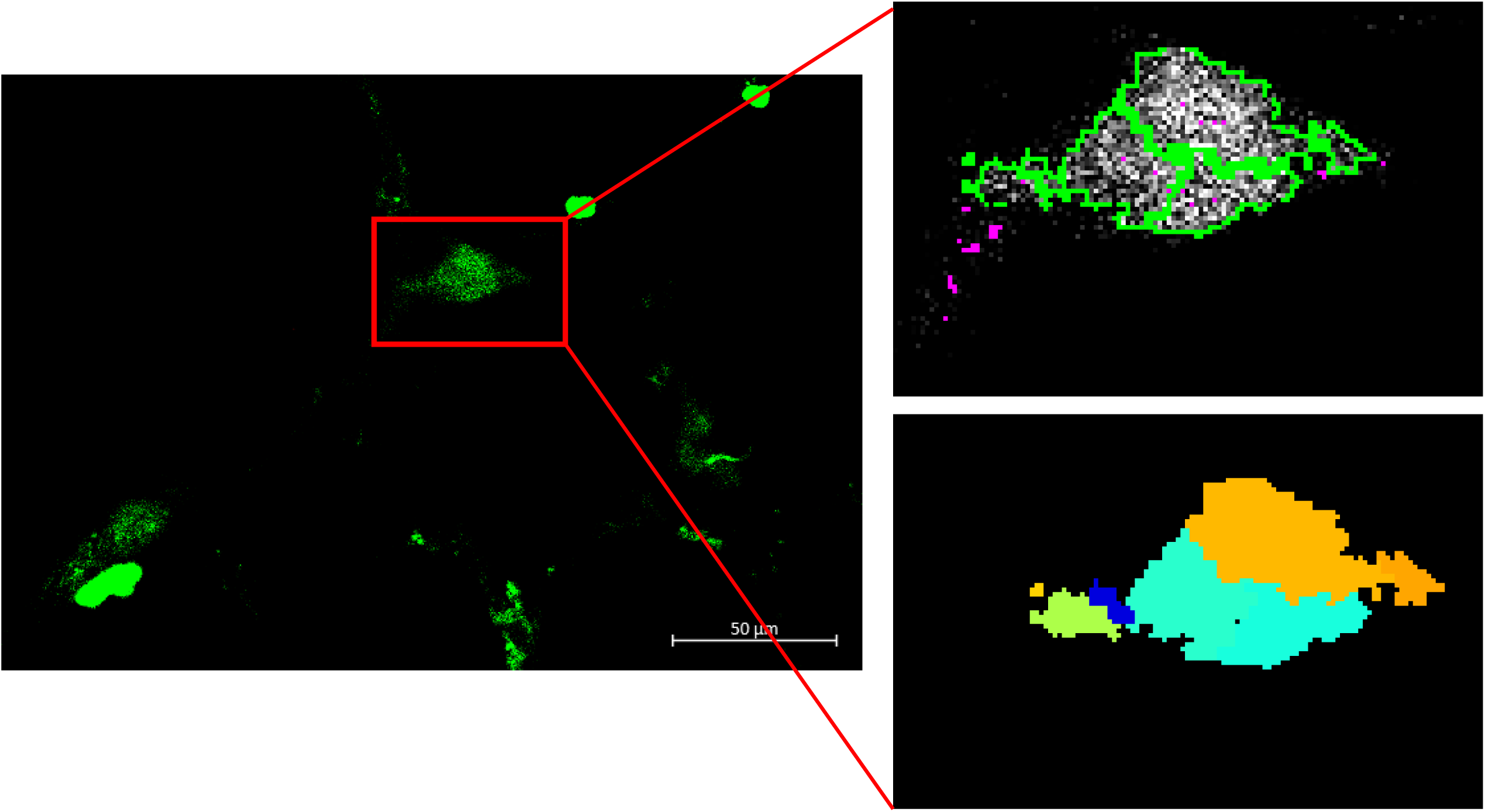
Analysis of Cell Clustering in immunostaining images. Representative image showing when multiple cells appeared clumped together were separated and counted, with the intensity of the signal used to distinguish individual cells.

**Figure. S11.**
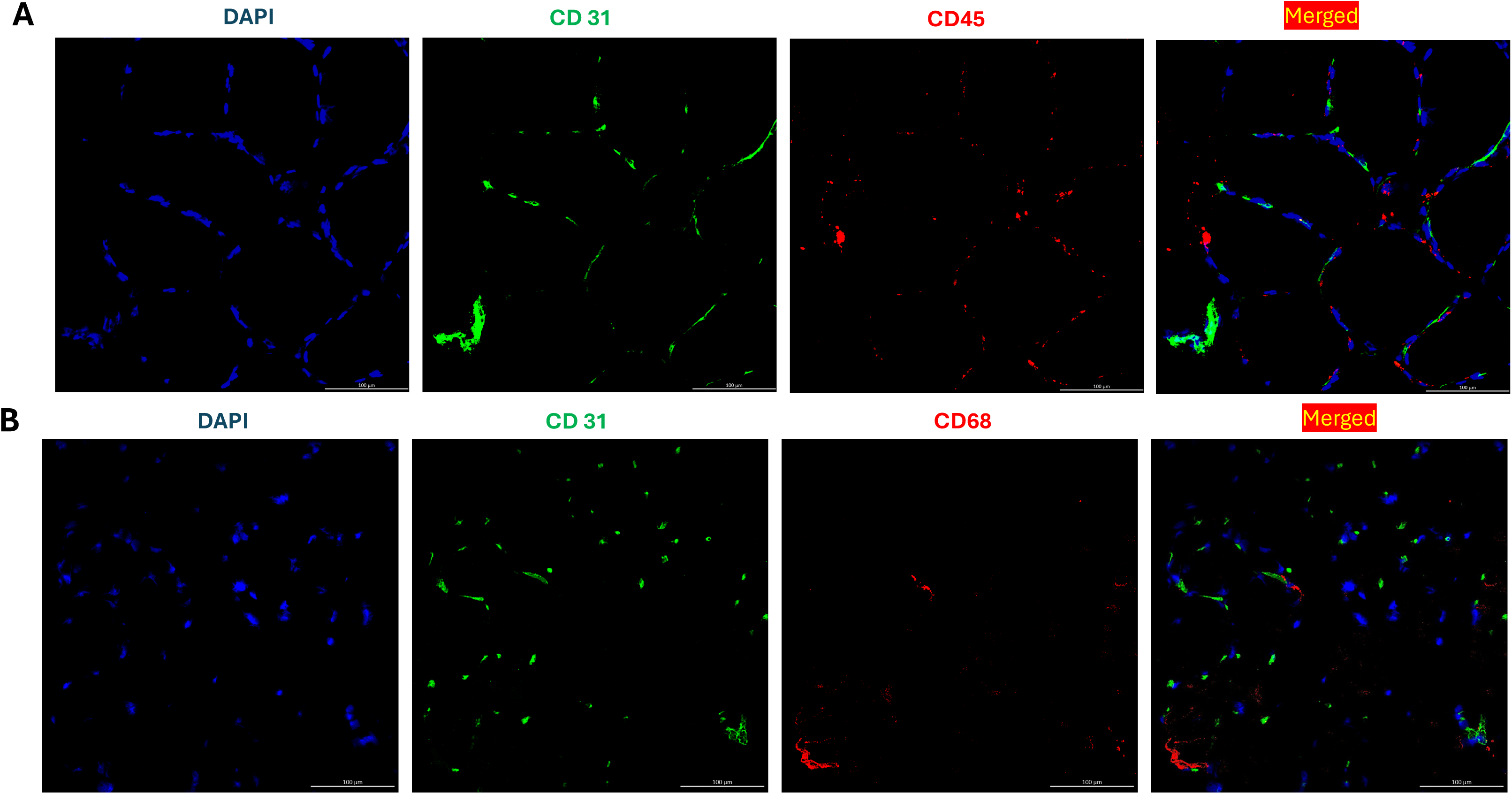
CD31 Immunostaining Specificity in Gastrocnemius Muscle Sections: Co-Staining with Immune Cell Markers. (A) DAPI (Blue), CD31 (EC marker) (Green) and CD45(Red) immunostaining was performed on gastrocnemius (GA) muscle sections from control (non-PAD) (scale bar: 100 *μ*m). Representative images were taken with 20X magnification. (**B**) DAPI (Blue), CD31 (EC marker) (Green) and CD68 “macrophage marker” (Red) immunostaining was performed on GA muscle sections from control (non-PAD) (scale bar: 100 *μ*m). Representative images were taken with 20X magnification.

